# Retinoic acid-mediated homeostatic plasticity drives cell type-specific CP-AMPAR accumulation in nucleus accumbens core and incubation of cocaine craving

**DOI:** 10.1101/2024.09.12.611703

**Authors:** Eun-Kyung Hwang, Amanda M. Wunsch, Marina E. Wolf

## Abstract

Incubation of cocaine craving, a translationally relevant model for the persistence of drug craving during abstinence, ultimately depends on strengthening of nucleus accumbens core (NAcc) synapses through synaptic insertion of homomeric GluA1 Ca^2+^-permeable AMPA receptors (CP-AMPARs). Here we tested the hypothesis that CP-AMPAR upregulation results from a form of homeostatic plasticity, previously characterized in vitro and in other brain regions, that depends on retinoic acid (RA) signaling in dendrites. Under normal conditions, ongoing synaptic transmission maintains intracellular Ca^2+^ at levels sufficient to suppress RA synthesis. Prolonged blockade of neuronal activity results in disinhibition of RA synthesis, leading to increased GluA1 translation and synaptic insertion of homomeric GluA1 CP-AMPARs. Using slice recordings, we found that increasing RA signaling in NAcc medium spiny neurons (MSN) from drug-naïve rats rapidly upregulates CP-AMPARs, and that this pathway is operative only in MSN expressing the D1 dopamine receptor. In MSN recorded from rats that have undergone incubation of craving, this effect of RA is occluded; instead, interruption of RA signaling in the slice normalizes the incubation-associated elevation of synaptic CP-AMPARs. Paralleling this in vitro finding, interruption of RA signaling in the NAcc of ‘incubated rats’ normalizes the incubation-associated elevation of cue-induced cocaine seeking. These results suggest that RA signaling becomes tonically active in the NAcc during cocaine withdrawal and, by maintaining elevated CP-AMPAR levels, contributes to the incubation of cocaine craving.

## INTRODUCTION

Relapse to cocaine use, which can occur even after long periods of abstinence, is a critical problem in treating cocaine use disorder. Relapse is often triggered by craving elicited by cocaine-associated cues ^1^. The persistence of vulnerability to cue-induced craving and relapse can be studied in rats using the ‘incubation of cocaine craving’ model ^2^. Incubation, which also occurs in abstinent human cocaine users ^3^, refers to the progressive intensification of cue-induced cocaine seeking that occurs over the first month of withdrawal from cocaine self-administration; seeking then remains high for months before declining ^2^.

Several brain areas contribute to incubation of cocaine craving ^4-6^. In the plateau phase (>30-40 d), we found that ‘incubated’ seeking depends on strengthening of AMPAR transmission onto medium spiny neurons (MSN) of the nucleus accumbens (NAc) core (NAcc) ^7, 8^. In drug-naive rats and in early withdrawal, this transmission is mainly mediated by AMPARs that contain the GluA2 subunit and are therefore Ca^2+^-impermeable or CI-AMPARs^7, 9^. AMPARs lacking the GluA2 subunit (Ca^2+^-permeable AMPARs, or CP-AMPARs) have higher conductance than CI-AMPARs, so their incorporation strengthens synapses ^10, 11^. Furthermore, by enabling Ca^2+^ entry at hyperpolarized potentials, CP-AMPARs affect induction of subsequent plasticity [e.g., ^12^]. Thus, CP-AMPAR plasticity has high functional significance. We observed that levels of homomeric GluA1 CP-AMPARs increase in NAcc MSN during incubation of cocaine craving, ultimately accounting for ∼25% of total AMPAR transmission ^9^. Once this occurs, blocking CP-AMPARs or removing them from NAcc synapses prevents expression of incubated cocaine craving ^9, 13, 14^.

It is important to understand the mechanisms that lead to CP-AMPAR upregulation and thus switch NAcc MSN into a state of persistently high cue-responsivity, as this may reveal novel targets for reducing craving. A clue is provided by the surprising observation that CP-AMPAR levels in NAcc rise only after ∼1 month of home-cage withdrawal ^7^, i.e., in the absence of any obvious Hebbian stimulus, leading us to propose that this represents homeostatic plasticity ^9^.

Synaptic scaling is a form of homeostatic plasticity that adjusts synaptic strength to compensate for changes in network activity. For example, prolonged inactivity can lead to strengthening of synapses ^15^. Synaptic scaling can utilize different induction and expression mechanisms in different cell types and under different conditions. Here we tested whether incubation of cocaine craving involves a form of homeostatic plasticity, originally discovered in hippocampal neurons, mediated by retinoic acid (RA) and expressed via synaptic insertion of homomeric GluA1 CP-AMPARs ^16^.

RA is a metabolite of retinol that is best known for activating transcription by binding to nuclear RA receptors (RARα, RARβ, and RARγ); this is critical for brain development and maintenance of the adult nervous system ^17, 18^. Transcriptional regulation in the NAc via RARs and related receptors (retinoic X receptors, or RXRs) is implicated in animal models of addiction ^19-22^. However, expression of RARα in cytoplasm increases during postnatal development ^23^ and it is present in hippocampal dendrites ^24, 25^. This dendritic RARα mediates a form of homeostatic synaptic plasticity that depends on regulation of translation rather than transcription ^16^. Briefly, under normal conditions, synaptic transmission maintains intracellular Ca^2+^ at levels sufficient to suppress dendritic RA synthesis through a mechanism involving calcineurin. Prolonged blockade of neuronal activity, by reducing intracellular Ca^2+^, disinhibits RA synthesis ^24, 26, 27^. RA then binds to dendritic RARα, rapidly leading to de-repression of GluA1 translation and synaptic insertion of homomeric GluA1 CP-AMPARs ^25, 28, 29^.

We hypothesize that this RA-dependent cascade is recapitulated in the NAcc during cocaine withdrawal. As detailed in the Discussion, a number of events occur in the NAcc during the period of withdrawal prior to CP-AMPAR upregulation that are consistent with reduced excitation of MSN; their cumulative effect may ultimately trigger RA synthesis. Here, using slice physiology and in vivo manipulations, we show that RA signaling becomes tonically active in the NAcc during cocaine withdrawal and that this is required for maintaining the elevated CP-AMPAR levels that mediate incubation of cocaine craving.

## MATERIALS AND METHODS

### Subjects

All procedures were approved by the Oregon Health & Science University Institutional Animal Care and Use Committee in accordance with the USPHS Guide for Care and Use of Laboratory Animals. We used wildtype (WT) and transgenic Long-Evans rats generated in our breeding colony; breeder rats for transgenic lines were obtained from the Rat Resource & Research Center and wildtype breeders from Charles River. To enable selective manipulation of the two populations of MSN in the NAc, we used previously validated knock-in rat lines encoding iCre recombinase immediately after the Drd1a or adenosine 2a receptor (Adora2a; A2a) loci, hereafter termed D1-Cre and A2a-Cre ^30^. A2a was targeted because it is selectively expressed in D2 MSN, while the D2 receptor is also expressed on cholinergic interneurons and other elements ^31^. In order to identify Cre+ neurons for slice recordings, D1-Cre or A2a-Cre rats were bred to reporter lines expressing either ZsGreen or TdTomato in the presence of Cre. Rats were allowed free access to food and water and maintained on a 12-h light/dark cycle. Rats were 10-15 weeks old at the onset of the experiment. More information on transgenic rats is provided in Supplementary Information.

### Surgery and cocaine self-administration

Rats destined for drug self-administration underwent jugular catheterization followed by extended-access intravenous self-administration (6 h/d x 10 d; each infusion paired with a light cue) of cocaine (0.5 mg/kg/infusion; NIDA Drug Supply Program) or saline (details in Supplementary Information).

### Cue-induced seeking tests

Some rats, destined for intra-NAcc infusion of vehicle or the RA synthesis inhibitor 4- (diethylamino)-benzaldehyde (DEAB) prior to cue-induced seeking tests, were implanted with both jugular catheters and bilateral intracranial guide cannulas aimed at NAcc (+1.4 A/P, ±2.35 M/L, -6.35 D/V) ^32, 33^ and received cue-induced seeking tests on withdrawal day (WD) 1 and WD50-60 following extended-access cocaine self-administration. The WD1 test was performed without manipulation to establish baseline craving. One hour prior to the WD50-60 test, the same rats received bilateral infusion (0.5 µl/side) of vehicle (0.1% DMSO in aCSF) or DEAB (50 µM in 0.1% DMSO in aCSF) into the NAcc. One hour after the injection, rats were placed in the self-administration chamber and the 1-h cue-induced seeking test was performed (see Supplementary Information).

### Slice electrophysiology

Electrophysiological experiments were performed as we have described previously (e.g., ^9, 13, 34-36^). Details are provided in Supplementary Information. Cocaine and saline rats were recorded between WD40 and WD60. All MSN were recorded from the NAcc subregion.

### Statistical Analyses

Clampfit (v11, Molecular Devices) and GraphPad Prism10 were used for figures and statistical analyses. All data were assessed for normality using Shapiro-Wilk tests. For comparison of two groups, Student’s t-tests (independent unless otherwise indicated) were used for normally distributed data and a Mann Whitney test was used for non-normally distributed data sets. Either one-way or two-way ANOVAs were utilized for comparing multiple groups followed by Tukey’s or Bonferroni post hoc multiple comparisons. For all analyses, significance was set at p<0.05. The data were presented as group mean ± S.E.M (see Supplementary Information).

## RESULTS

### RA increases synaptic CP-AMPAR levels in NAcc MSN from drug-naïve rats

Our overarching hypothesis is that RA synthesis is suppressed in NAcc MSN of drug-naïve rats, but reductions in neuronal activation during forced abstinence from extended-access cocaine self-administration lead to RA-dependent homeostatic upregulation of CP-AMPARs. A prediction of this hypothesis is that increasing RA signaling in MSN of drug-naïve rats should increase synaptic CP-AMPAR levels while inhibiting RA signaling should have no effect. We tested this using whole cell patch clamp recordings in NAcc MSN from drug-naïve male and female rats.

First, slices from drug-naïve rats were preincubated with RA (2 µM) or vehicle (0.01% DMSO) for 1 h prior to recordings. We generated I-V curves for the AMPAR-mediated evoked EPSC (eEPSC) to assess inward rectification, a hallmark of CP-AMPARs. While MSN from vehicle-preincubated slices exhibited a linear I-V relationship, as expected from our prior results (e.g., ^9^), RA-preincubated slices showed inward rectification. A two-way ANOVA with treatment (Veh or RA) as a between-subject factor and voltage (-80, -60, -40, -20, 0, 20, 40 mV) as a within-subject factor showed a main effect of treatment (F_(1, 32)_=5.772, p=0.0223) and holding potential (F_(3.001, 96.04)_=3448, p<0.0001), and a treatment x holding potential interaction (F_(6, 192)_=8.197, p<0.0001) (Fig. 1A, B), which was quantified as the rectification index or RI [eEPSC_-70mV_/(-70- E_rev_)]/[eEPSC_+40mV_/(+40-E_rev_)]. The RI was significantly higher in MSN from RA-treated slices than vehicle-treated slices (Fig. 1C, unpaired t-test, t_(32)_=5.509, p<0.0001). To confirm these results, eEPSCs were measured before and after bath application of the selective CP-AMPAR antagonist 1-Naphthylacetyl spermine trihydrochloride (NASPM; 100 µM). MSN preincubated with RA displayed a significant NASPM-induced reduction in eEPSC amplitude, whereas vehicle-preincubated MSN did not (Fig. 1D-E, two-way ANOVA, time x drug history interaction, F_(75,1350)_=3.093, p<0.0001; main effect of time, F_(10.30, 185.5)_=2.410, p=0.0095; main effect of drug history, F_(1, 18)_=38.79, p<0.0001). Mean eEPSC amplitudes, measured 15-20 min after NASPM application and expressed as percent of baseline, were significantly reduced in RA-pretreated MSN (Fig. 1F, unpaired t-test, t_(18)_=6.431, p<0.0001). To confirm results obtained after RA pretreatment, slices from drug-naïve rats were preincubated for 1-2 h with a selective agonist of RARα, AM580 (5 µM), or vehicle (0.02% DMSO). As observed after RA, AM580 produced inward rectification (Fig. 1G, two-way ANOVA, Vh x treatment interaction, F_(6, 150)_=9.253, p<0.0001; main effect of Vh, F_(3.229,80.73)_=1810, p<0.0001; main effect of treatment, F_(1, 25)_=4.710, p=0.0397) and an elevated RI (Fig. 1H; unpaired t-test, t_(25)_=2.821, p=0.0092). Furthermore, preincubation of slices with RA in the presence of a selective antagonist of RARα, Ro 41-5253, prevented RA-induced inward rectification. Ro 41-5253 pretreatment had no effects on the current-voltage relationship of AMPAR-mediated eEPSCs (Fig 1I-J, unpaired t-test, t_(39)_=0.2016, p=0.8413).

**Figure 1.**
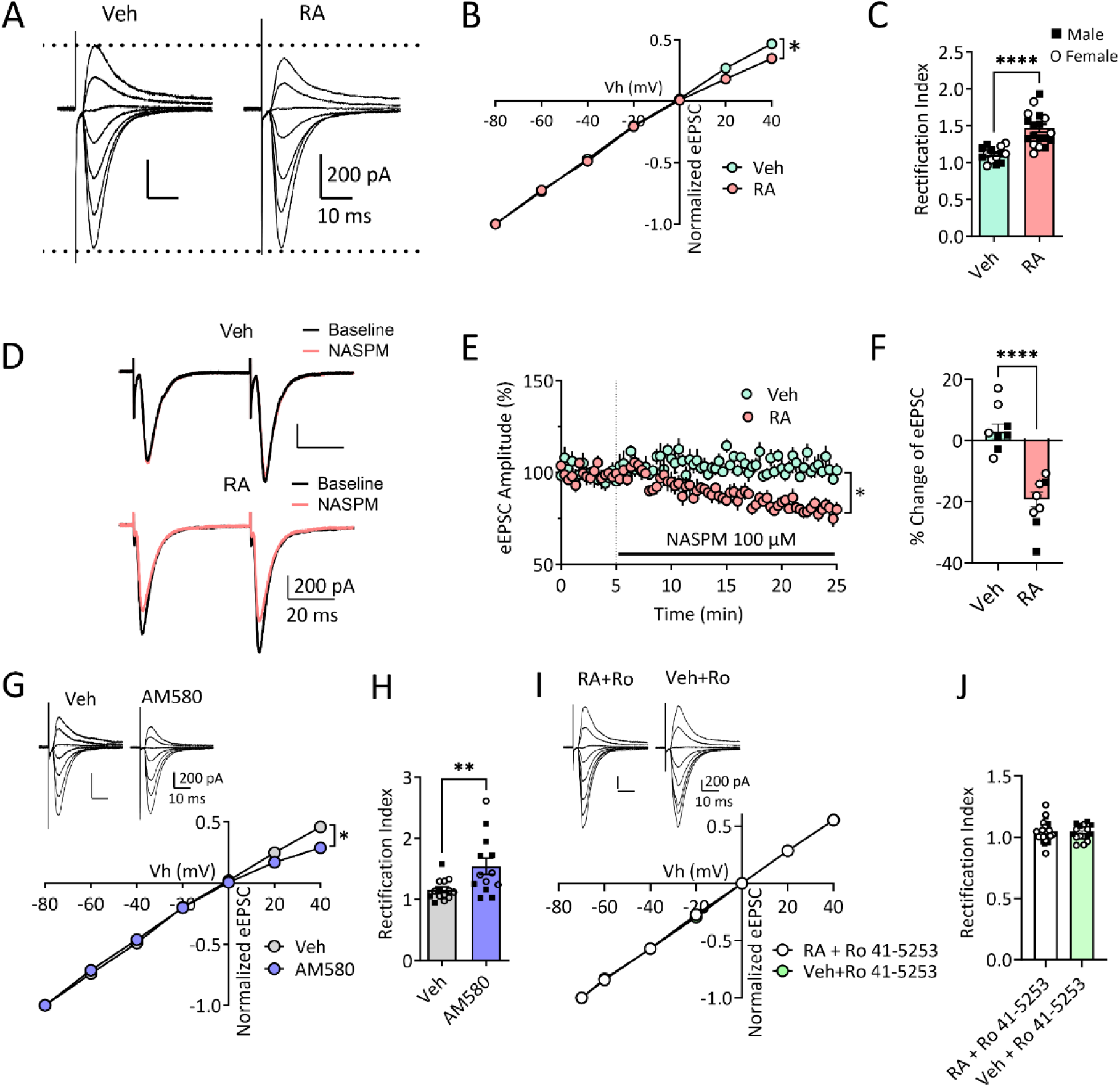
Retinoic acid (RA) induces upregulation of synaptic CP-AMPARs in nucleus accumbens core (NAcc) medium spiny neurons (MSN). A, AMPAR-mediated eEPSCs recorded at -80, -60, -40, -20, 0, 20, 40 mV from NAcc MSN in drug-naïve rats after pretreatment with vehicle (Veh, 0.01% DMSO) or RA (2 µM) for 1-2 h. Scale bar: 200 pA/10 ms. B, I-V plots of normalized AMPAR-mediated synaptic responses at the different membrane holding potentials (Vh). The MSN from Veh-pretreated slices showed a linear I-V relationship, whereas inward rectification was detected in MSN after RA pretreatment [two-way ANOVA, Sidak’s multiple comparisons test, Veh vs RA at Vh=20mV, p=0.0077; at Vh=40mV, p=0.0015; Veh, 13 cells/7rats (3M, 4F); RA, 18 cells/10 rats (5M, 5F)]. * indicates significant main effects of treatment (p=0.0223) and Vh (p<0.0001), as well as Vh x treatment interaction (p<0.0001). C, Quantification of rectification index (RI) showing a significantly higher RI after pretreatment with RA compared to Veh [unpaired t-test, p<0.0001; Veh, 13 cells/7 rats (3M, 4F); RA, 18 cells/10 rats (5M, 5F)]. M, male; F, female. D, Representative eEPSC traces (black line, averaged baseline, 0-5 min; red line, 15-20 min after NASPM application). Scale bar: 200 pA/20 ms. E, Time course before and during 100 µM NASPM application 1 h after pretreatment of slices with Veh or RA. * indicates significant main effects of treatment and time, as well as Vh x time interaction (all p<0.0001). F, % change of mean eEPSC amplitudes measured 15-20 min after NASPM application [unpaired t-test, p<0.0001; Veh 9 cells/6 rats (3M, 3F); RA, 11 cells/5 rats (3M, 2F)]. G, Example traces of AMPAR-mediated eEPSCs at -80, -60, -40, -20, 0, +20 and +40 mV recorded from MSN after Veh or AM580 pretreatment (1-2 h). Scale bar: 200 pA/10 ms. I-V plots of normalized AMPAR-mediated synaptic responses of MSN after Veh or AM580 (5 µM) pretreatment. Veh-pretreated MSN showed a linear I-V relationship, while an inward rectifying I-V relationship was observed in MSN after pretreatment with AM580 (two-way ANOVA, Sidak’s multiple comparisons test, Veh vs. AM580 at 20 mV, p=0.677; at 40 mV, p<0.0001). * indicates significant main effects of treatment (p=0.0397) and Vh (p<0.0001), as well as Vh x treatment interaction (p<0.0001). H, The RI of AM580-pretreated MSN was significantly higher compared to Veh-pretreated MSN [unpaired t-test, **p=0.0092; Veh, 13 cells/5 rats (3M, 2F); AM580, 14 cells/5 rats (2M, 3F)]. I, Example traces of eEPSCs recorded at membrane potentials from -70 mV to +40 mV after pretreatment with RA or Veh (1-2 h) in the presence of Ro 41-5253, a selective antagonist of RARα. eEPSCs are normalized to peak amplitude at -70 mV. Scale bars, 200pA/10ms. The I-V plots of normalized eEPSCs recorded after pretreatment with either Veh or RA in the presence of Ro 41-5253 showed linear relationships (two-way ANOVA, p>0.05). J, Ro 41-5253 blocked RA-induced elevation of RI. Ro 41-5253 (+Veh) had no effects on RI [unpaired t-test, p>0.05; RA+Ro 41-5253, 25 cells/ 6 rats (2M,4F); Veh+Ro 41-5253, 16 cells/4 rats (2M, 2F)]. Data are presented as mean ± S.E.M. Individual data points for each rat are included in bar graphs (▪ Male, O Female).

We also determined whether activation of RA signaling affects other properties of NAcc MSN from drug-naïve rats. Consistent with the higher single channel conductance of CP-AMPARs^10, 11^, we observed an increase in sEPSC amplitude in RA-pretreated MSN compared to vehicle-preincubated MSN [Fig. 2A-B; unpaired t test, t_(22)_=4.578, p=0.0001]. However, RA preincubation did not affect sEPSC frequency (Fig. 2C, unpaired t-test, t_(22)_=0.8084, p=0.4275) or the paired-pulse ratio (PPR), suggesting no change in presynaptic function (Fig. 2D, two-way ANOVA; ISI x treatment interaction, F_(5,80)_=0.5606, p=0.7298; main effect of treatment, F_(1,16)_=0.3862, p=0.5431). RA also failed to alter passive membrane properties (Fig. 2E, resting membrane potential: unpaired t test, t_(44)_=0.2191, p=0.8276; Fig. 2F, input resistance: unpaired t test, t_(46)_=0.1002, p=0.9206; Fig. 2G, I-V: two-way ANOVA, injected current x treatment interaction, F_(10, 460)_=0.08008, p>0.999, main effect of treatment, F_(1,46)_=1.804, p=0.1858) or active membrane properties (Fig. 2H, excitability: two-way ANOVA, injected current x treatment interaction; F_(12,516)_=1.234, p=0.2558, main effect of treatment, F_(1, 43)_=2.257, p=0.1404; Fig. 2I, rheobase current: unpaired t test, t_(44)_=0.006727, p=0.9947) of NAcc MSN from drug-naïve rats.

**Figure 2.**
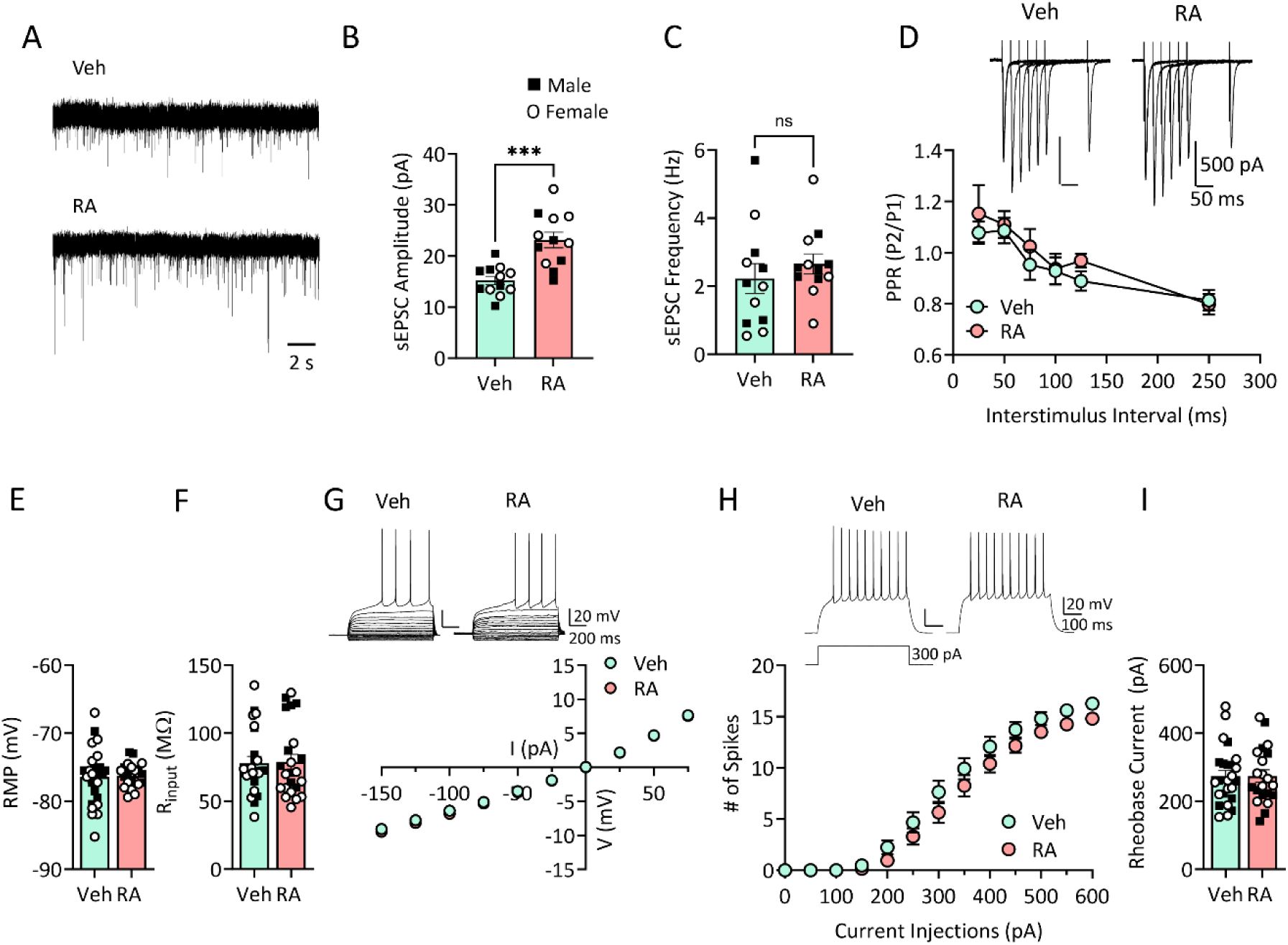
Retinoic acid (RA) causes postsynaptic alterations in nucleus accumbens core (NAcc) medium spiny neurons (MSN) without affecting presynaptic functions or intrinsic membrane properties. A, Representative traces of spontaneous EPSCs (sEPSC) recorded from NAcc MSN in slices pretreated (1-2 h) with either vehicle (Veh, 0.01 % DMSO) or RA (2 µM). Scale bar, 2s. B-C, Quantification of sEPSC amplitude (B) and frequency (C) after pretreatment with Veh or RA. sEPSC amplitude (unpaired t-test, ***p<0.01), but not frequency (unpaired t-test, p=0.4275), was increased in MSN after RA pretreatment [Veh: 12 cells/5 rats (3M, 2F); RA: 12 cells/5 rats (3M, 2F)]. ns, not significant. D, Mean paired-pulse ratios of AMPAR-mediated eEPSCs at different inter stimulus intervals (ISIs) in NAcc MSN after pretreatment with Veh or RA. No difference between Veh and RA-treated group [(two-way ANOVA, Sidak’s multiple comparisons test, p>0.05; Veh: 9 cells/6 rats (3M, 3F); RA: 9 cells/5 rats (3M, 2F)], indicating that presynaptic release probability is not changed after RA treatment. Top: Representative traces from individual cells in each group. E, Average resting membrane potential (RMP) did not differ between Veh- and RA-treated NAcc MSN [unpaired t-test, p=0.8276; Veh: 25 cells/6 rats (3M, 3F); RA: 21 cells/6 rats (3M, 3F)]. F-G, Mean input resistance was unchanged after RA pretreatment (unpaired t-test, p=0.9206). Representative traces and current-voltage (I-V) plots recorded with incremental subthreshold current injections (step current injections, -150 pA-75 pA/Δ25 pA, 1s) [two-way ANOVA, Sidak’s multiple comparisons test, p>0.05; Veh: 25 cells/6 rats (3M, 3F); RA: 23 cells/6 rats (3M, 3F)]. H, Representative traces (top) of action potentials in response to 300 pA current injections in NAcc MSN after Veh or RA pretreatment. Mean firing versus current relationship (excitability), measured as the number of action potentials as a function of current injected (0-600 pA, Δ50 pA, 500 ms duration), did not differ between Veh and RA-treated MSN [two-way ANOVA, Sidak’s multiple comparisons test, p>0.05; Veh: 25 cells/6 rats (3M, 3F); RA: 21 cells/6 rats (3M, 3F)]. I, Rheobase current was not changed after RA pretreatment compared to Veh [unpaired t-test, p=0.9947; Veh: 25 cells/6 rats (3M, 3F); RA: 21 cells/6 rats (3M, 3F)].

### RA signaling is tonically active in NAcc MSN after incubation of cocaine craving and is required to maintain elevated CP-AMPAR levels

Whereas RA signaling appears to be suppressed in NAcc MSN of drug-naïve rats (Fig. 1), our overarching hypothesis predicts tonic activation of RA signaling in NAcc MSN from rats that have undergone incubation of cocaine craving. This in turn predicts that increasing RA signaling will have no effect on CP-AMPAR levels in these MSN (occlusion) whereas inhibiting RA signaling may reduce CP-AMPAR levels. To test these predictions, we assessed NAcc MSN following extended-access saline or cocaine self-administration [Fig. 3A-E; see legend to Fig. 3B for statistical analysis]. Recordings were performed between WD40 and WD60, a period when incubation and CP-AMPAR levels have plateaued in cocaine rats ^7^. In MSN from saline rats, preincubation with RA elevated the RI, as expected from parallel studies in drug-naïve rats (Fig. 1), but preincubation with the RA synthesis inhibitor DEAB (0.05% DMSO) had no effect (Fig. 3G, two-way ANOVA, Vh x treatment interaction, F_(16,256)_=15.82, p<0.0001; main effect of Vh, F_(1.719, 55.02)_=6229, p<0.0001; main effect of treatment, F_(2, 32)_=9.678, p=0.0005; Tukey’s multiple comparisons test, Veh vs RA, p<0.0001; Veh vs DEAB, p=0.9961; RA vs DEAB, p<0.0001; Fig. 3H, one-way ANOVA, F_(2,32)_=18.76, p<0.0001). Comparing vehicle preincubated slices from saline and cocaine rats, the RI was elevated in MSN from cocaine rats compared to the saline group (t-test, p<0.0001), consistent with our prior results showing an elevated RI after incubation of cocaine craving (see Introduction). Importantly, RA preincubation produced no further elevation of the RI in cocaine MSN, whereas DEAB reduced the RI relative to vehicle (Fig. 3J, two-way ANOVA, Vh x treatment interaction, F_(16,296)_=16.06, p<0.0001; main effect of Vh, F_(1.849, 68.40)_=8145, p<0.0001; main effect of treatment, F_(2, 37)_=29.80, p<0.0001; Fig. 3K, one-way ANOVA, F_(2,38)_=20.26, p<0.0001; Tukey’s multiple comparisons test, Veh vs. RA, p=0.9428; Veh vs. DEAB, p<0.0001; RA vs. DEAB, p<0.0001). Neither cocaine nor saline MSN showed evidence of presynaptic changes after RA or DEAB compared to vehicle treatment (Fig. S1). Together with Fig. 1, these results indicate that RA signaling is not occurring in saline rats but is tonically activated in the NAcc of cocaine rats.

**Figure 3.**
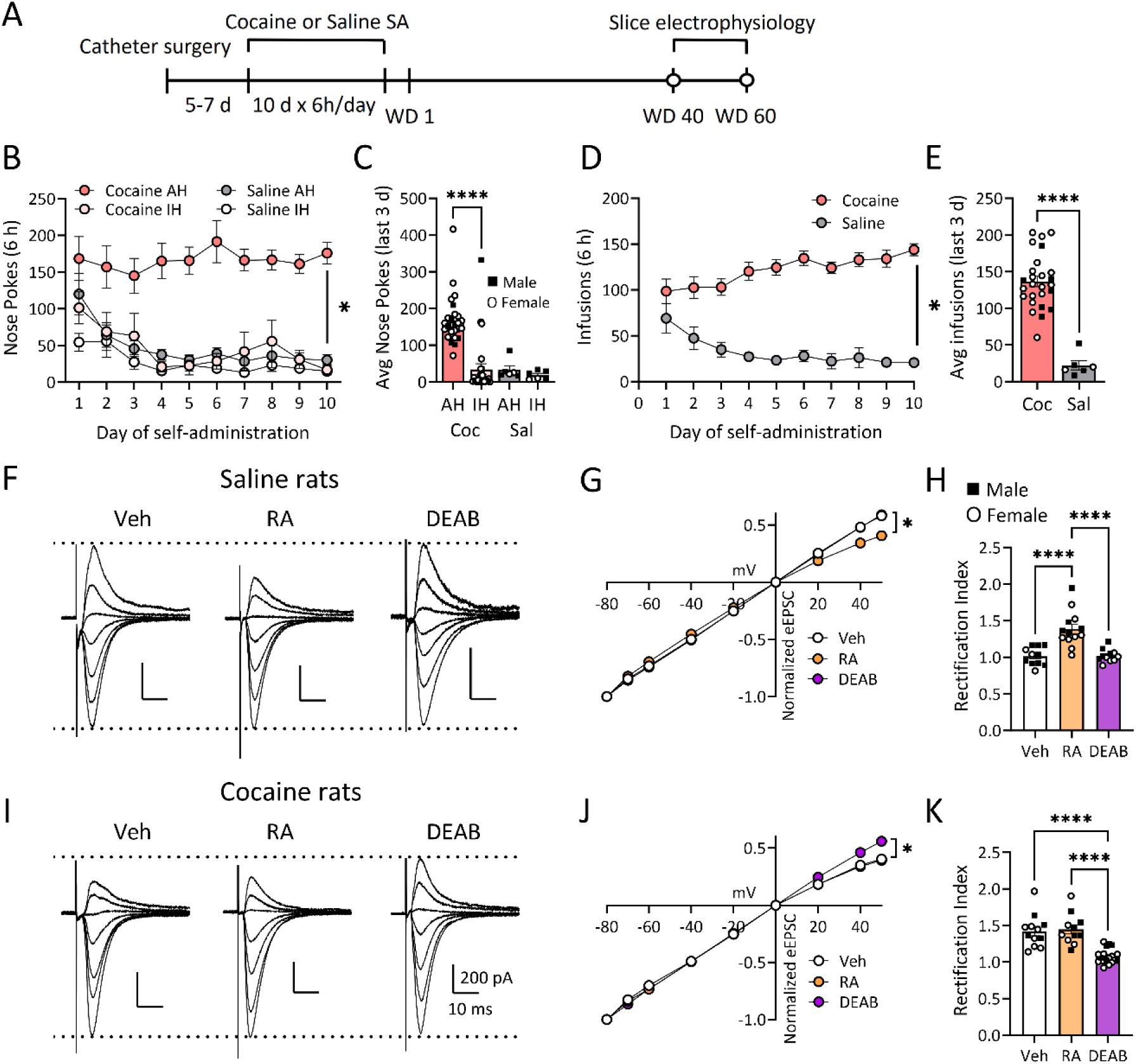
Pretreatment with the retinoic acid (RA) synthesis inhibitor DEAB blocks RA- or cocaine-mediated upregulation of synaptic CP-AMPARs in nucleus accumbens core (NAcc) medium spiny neurons (MSN). A, Experimental timeline. Male (M) and female (F) rats underwent 10 days of saline (Sal) or cocaine (0.5 mg/kg/infusion) self-administration (SA) training (6 h/day) followed by a drug-free abstinence period. On withdrawal day (WD) 40-60 rats were used for slice recordings. B, Training data for rats used in electrophysiology experiments (includes all cocaine and saline rats used for Figs. 3 & 5; see Fig. S2 for data disaggregated for wild-type and transgenic rats). Pokes in the active hole (AH) resulted in an infusion of cocaine paired with a light cue. The average number of AH nose-pokes is higher than inactive hole (IH) nose-pokes throughout 10 days of cocaine SA [two-way ANOVA, day x drug interaction, F_(27,540)_=0.8576, p=0.6746; main effect of day, F_(3.072,184.3)_=0.1840, p=0.1840; main effect of drug, F_(3,60)_=19.95, p<0.0001; Sidak’s multiple comparisons post hoc test: Coc AH vs Coc IH, from day 4, p<0.0001; Coc, 26 rats (7M, 19F), Sal, 6 rats (3M, 3F)]. * indicates significant main effects of drug (p<0.0001). C, Cocaine rats showed a significantly higher average number of AH nose-pokes in the last 3 days of SA compared to IH nose pokes (two-way ANOVA, hole type x drug interaction, F_(1,60)_=8.288, p=0.0055; main effect of hole type, F_(1,60)_=12.52, p=0.0008; main effect of drug, F_(1,60)_=13.17, p=0.0006; Tukey’s multiple comparisons test: ****p<0.0001, Coc AH vs. Coc IH; p= 0.0001, Coc AH vs. Sal AH). There was no difference between AH and IH nose-pokes in Sal rats. D, The number of Coc or Sal infusions taken over the 10 days of SA training [two-way ANOVA, day x drug interaction, F_(9,270)_=4.329, p<0.0001; main effect of day, F_(2.253, 67.58)_=0.4471, p=0.6646; main effect of drug, F_(1,30)_=36.83, p<0.0001; Sidak’s multiple comparisons test: p=0.0002, Coc vs. Sal from SA day 3; p<0.0001, Coc vs. Sal from SA day 4-10; Coc, 26 rats (7M, 19F), Sal, 6 rats (3M, 3F)). * indicates significant main effects of drug (p<0.0001). E, The average number of Coc infusions in the last 3 days of SA training is significantly higher than Sal infusions (unpaired t-test, t_(30)_=7.386, ****p<0.0001). F, Representative traces of AMPAR-mediated synaptic responses recorded from Sal rats (WD 40-60) at -80, -60, -40, -20, 0, +20 and +40 mV, normalized to peak amplitude at -80 mV. Scale bar: 200 pA/10ms. G, I-V plot of normalized eEPSCs at different holding potentials for NAcc MSN from Sal rats after preincubation with Veh, RA or DEAB. The RA-preincubated MSN showed inward rectification of I-V relationship, while Veh-pretreated MSN showed a linear I-V relationship. Co-treatment of DEAB with RA reversed RA effects (two-way ANOVA, Sidak’s multiple comparisons test: Veh vs RA at Vh=20mV, p=0.0062; at Vh=40mV, p=0.0002; at Vh=50mV, p=0.0003; RA vs DEAB at Vh=20mV, p=0.0004, at Vh=40mV and 50 mV, p<0.0001). * indicates significant main effects of treatment and Vh, as well as Vh x treatment interaction (all p<0.0001). H, Quantification of mean rectification index (RI) showing a higher RI after RA pretreatment compared to Veh or DEAB pretreatment in saline rats [one-way ANOVA, Tukey’s multiple comparison tests: ****p<0.0001, Veh vs. RA; ****p<0.0001, RA vs. DEAB; Veh, 11 cells/4 rats (2M, 2F); RA, 13 cells/4 rats (2M, 2F); DEAB: 11 cells/4 rats (2M, 2F). I-K, Representative traces (I) of AMPAR-mediated EPSCs from Coc rats (WD40-60) at -70 to +40 mV in NAcc MSN. AMPAR I-V curves [J, two-way ANOVA, Sidak’s multiple comparisons test: Veh vs DEAB at Vh=20-50mV, p<0.0001; RA vs DEAB at Vh=20-50mV, p<0.0001; * indicates significant main effects of treatment and Vh, as well as Vh x treatment interaction (all p<0.0001)] and RI of MSN after Veh, RA or DEAB pretreatment [K, one-way ANOVA, Tukey’s multiple comparison tests: Veh vs DEAB, ****p<0.0001; RA vs DEAB, ****p<0.0001; Veh, 12 cells/ 5 rats (2M, 3F); RA, 11 cells/5 rats (2M, 3F); DEAB, 18 cells/6 rats (2M, 4F)]. Data are presented as mean ± S.E.M. Individual data points for each rat are included in bar graphs (▪ Male, O Female).

### Blocking RA synthesis prior to a seeking test prevents expression of incubated craving

As CP-AMPAR activation in the NAcc is required for expression of cocaine incubation ^9, 13, 35^ and 1-2 h of preincubation with DEAB removes CP-AMPARs from NAcc synapses (Fig. 3), we reasoned that intra-NAcc DEAB infusion might prevent the expression of cocaine incubation. To test this, rats were implanted with bilateral intracranial guide cannulas at the time of jugular catheterization and trained to self-administer cocaine (Fig. 4A-D). They received seeking tests on WD1 (to assess baseline craving) and WD50-60 (when incubation is maximal). Rats destined to receive vehicle and DEAB took a similar number of cocaine infusions over the 10 days of cocaine self-administration [Fig. 4D, two-way ANOVA with treatment (between-subject) and withdrawal day (within-subject) as factors; no significant treatment X WD interaction or main effect of treatment, p>0.05]. Matching the timing of electrophysiological experiments, vehicle or DEAB was bilaterally infused into NAcc through a guide cannula 1 h prior to the WD50-60 seeking test (Fig. 4E). Whereas vehicle-infused rats expressed incubation on WD50-60 as expected, rats receiving intra-NAcc DEAB did not express incubation (Fig. 4E, two-way ANOVA with treatment (between-subject) and WD (within-subject) as factors; treatment x WD interaction F_(1,14)_=6.068, p=0.0273; main effect of treatment F_(1,14)_=3.118, p=0.0629; post hoc Sidak’s multiple comparisons test, p=0.0129, WD1 vs. WD50-60 for vehicle-infused rats; p=0.008, WD50-60 vehicle vs DEAB). The incubation score (WD50-WD1 active nose pokes), a measure of the magnitude of incubation of cocaine craving, was significantly lower in DEAB-infused rats compared to vehicle-infused rats (Fig. 4F, Mann-Whitney, U=8, p=0.0115). Rats that received intra-NAcc DEAB infusion showed a similar number of inactive pokes and beam breaks during the WD50-60 seeking test compared to vehicle-infused control rats (Fig. 4G-H), suggesting that DEAB infusions into the NAcc attenuated expression of incubated cocaine craving without altering general locomotor activity or discrimination between nose poke holes. These results indicate that tonic RA signaling after prolonged cocaine withdrawal is required for maintenance of CP-AMPARs which in turn enable expression of incubation.

**Figure 4.**
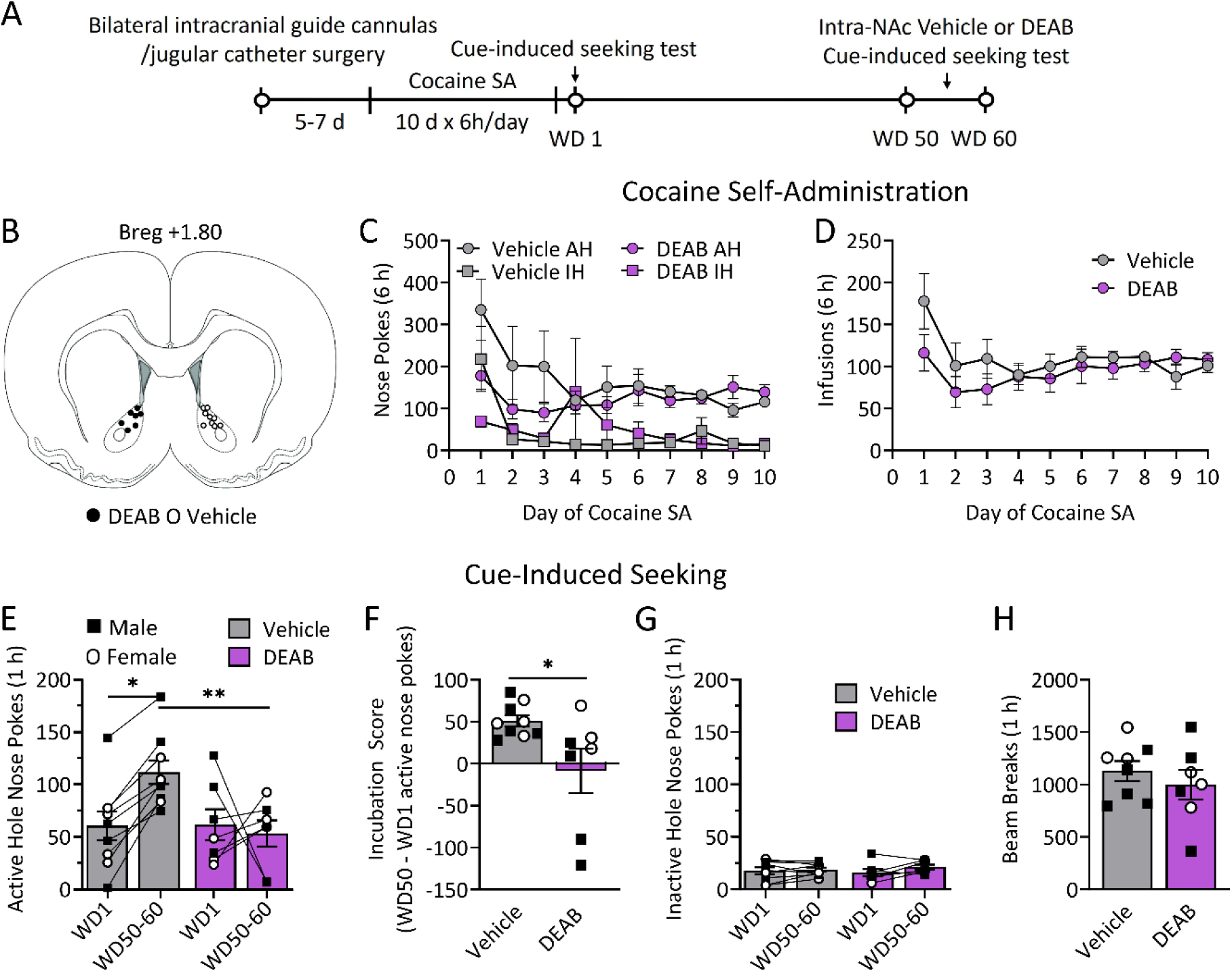
Acute inhibition of retinoic acid (RA) synthesis in the nucleus accumbens core (NAcc) with DEAB reduces the expression of incubation of cocaine craving. A, Experimental timeline of cocaine self-administration (SA) training (6 h/day for 10 days) and cue-induced seeking tests with DEAB or vehicle microinjections (DEAB, 50 µM/0.5 µl/side; Vehicle, 0.1% DMSO) administered into NAcc 1 h before the seeking test. All rats underwent two seeking tests (WD1 and WD50-60) and received vehicle or DEAB infusions on WD50-60 (no manipulation on WD1). B, Approximate placement of cannula tips in the NAcc (bregma +1.80). C, Cocaine SA training data for male (9 rats) and female (7 rats) destined to receive intra-NAcc infusion of vehicle or DEAB prior to the WD50-60 seeking test. All rats learned to discriminate between the active hole (AH) and inactive hole (IH) [two-way ANOVA with treatment (between subject) and SA day (within subject) as factors; treatment x day interaction F_(27,14)_=1.501, p=0.0586; main effect of treatment F_(3,27)_=6.175, p=0.0025; post hoc Sidak’s multiple comparisons test: SA days 4-6, p<0.05 and SA days 7-10, p<0.001, AH vs IH for rats destined for vehicle; SA days 7-10, p<0.001, AH vs IH for rats destined for DEAB]. D, Rats destined for intra-NAcc infusion of vehicle or DEAB took similar numbers of cocaine infusions over the 10 days of SA training. E, Infusion of DEAB into the NAcc 1 h prior to a seeking test on WD50-60 significantly reduced cocaine seeking compared to vehicle-infused controls [two-way Mixed ANOVA, Sidak’s multiple comparisons post hoc test: *p=0.0129, vehicle WD1 vs. WD50-60; **p=0.008, vehicle WD50-60 vs. DEAB WD50-60; p>0.05, DEAB WD1 vs. WD50-60; Veh, 9 rats (5M, 4F); DEAB, 7 rats (4M, 3F)]. F, The incubation score (WD50 - WD1 active nose-pokes) was significantly lower in DEAB infused rats compared to vehicle infused rats (Mann Whitney test, U=8 *p=0.0115). G, Inactive nose pokes were similar between vehicle and DEAB infused rats on WD1 and WD50-60 (p>0.05). H, The number of beam breaks (a measure of locomotor activity) during the WD50-60 cue-induced seeking test was similar between vehicle and DEAB infused rats (p>0.05). Data are presented as mean ± S.E.M. Individual data points for each rat are included in bar graphs (▪ Male, O Female).

### RA-dependent homeostatic plasticity occurs selectively in D1 MSN

Based on several lines of evidence (see Discussion), we predicted that the CP-AMPAR upregulation which underpins incubation is occurring in D1 MSN and therefore RA signaling is preferentially occurring in D1 MSN. To visualize D1 MSN for slice recordings, we crossed D1-Cre rats with Cre reporter lines that express ZsGreen or TdTomato. On WD40-60 after saline or cocaine self-administration (see Fig. S2 for self-administration data), we recorded AMPAR eEPSCs in D1+ (fluorescent) MSN or D1-MSN (presumed D2 MSN) (Fig. 5A, B). We observed a linear IV relationship in D1+ MSN from saline rats, as expected, whereas in cocaine rats inward rectification was observed in D1+ MSN but not D1- MSN (Fig. 5C, two-way RM ANOVA, Vh x cell type/drug history interaction, F_(18,456)_=12.14, p<0.0001; main effect of Vh, F_(2.497,189.8)_=178966, p<0.0001; main effect of cell type/drug history, F_(3,75\)_=19.88, p<0.0001; Fig. 5D, one-way ANOVA, F_(3,90)_=39.20, p<0.0001; Tukey’s multiple comparisons test, D1+ Sal vs. D1+ Coc, p<0.0001; D1+ Coc vs. D1- Coc, p<0.0001; D1+ Coc vs. D1- Sal, p<0.0001]. Confirming these results, the CP- AMPAR-selective antagonist NASPM selectively reduced eEPSC amplitude only in D1+ MSN from cocaine rats (Fig. 5E, two-way ANOVA, time x cell-type/drug history interaction, F_(120,1500)_=3.401, p<0.0001; main effect of time, F_(10.07,251.8)_=2.835, p=0.0023; main effect of cell-type/drug history, F_(2,25)_=16.01, p<0.0001; Fig. 5F, one-way ANOVA, F_(2,25)_=16.04, P<0.0001; Tukey’s multiple comparisons, D1+ Sal vs. D1+ Coc, p=0.0003; D1+ Sal vs. D1- Coc, p=0.8204; D1+ Coc vs. D1- Coc, p=0.0002). Extending these results, we found increased excitatory tone in D1 MSN from cocaine rats compared to saline rats (increased EI balance, increased eEPSC amplitude in response to electrical stimulation, increased excitability, and decreased rheobase; Fig. S3). To confirm results from D1- MSN (presumed D2 MSN), we used A2a-Cre transgenic rats (the A2a receptor colocalizes with the D2 receptor in MSN; see Methods) crossed with A2a ZsGreen or TdTomato reporter rats. A2a+ MSN from cocaine rats demonstrated a linear IV and no NASPM sensitivity (Fig. S4A-D), as well as no change in excitatory tone (Fig. S4E-K).

**Figure 5.**
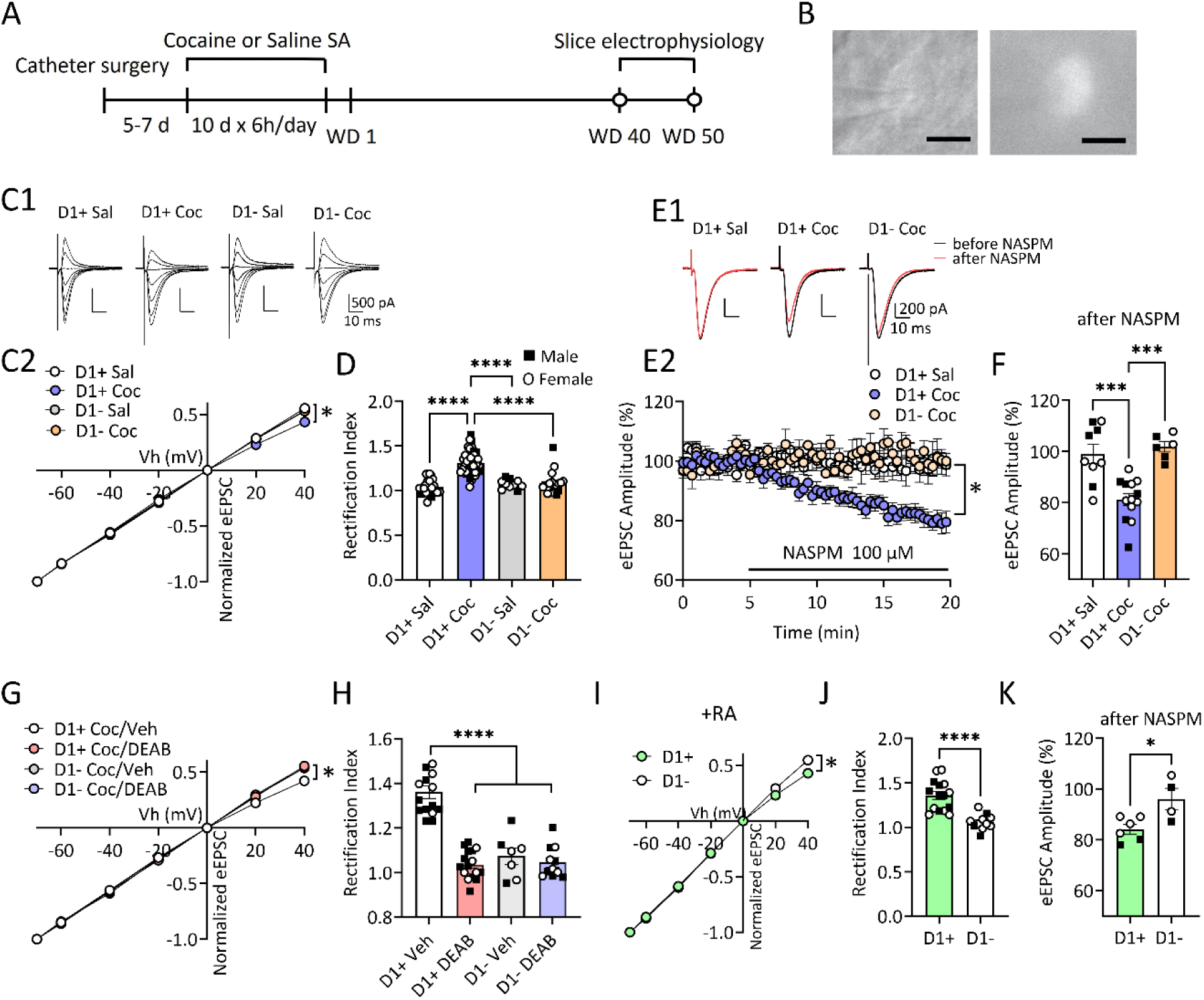
CP-AMPAR upregulation via retinoic acid (RA) signaling occurs preferentially in D1+ medium spiny neurons (MSN) in the nucleus accumbens core (NAcc). A, Experimental timeline. D1-Cre rats (from crosses with TdTomato or ZsGreen reporter lines) underwent 10 days of saline (Sal) or cocaine (Coc) self-administration (SA) training followed by a drug-free abstinence period (C-H). Slice recordings were done on withdrawal day (WD) 40-60. B, Representative images showing fluorescence-expressing (D1 positive) MSN in NAcc in a slice from a D1-tdTomato rat. Bright field image at 40x magnification. Scale bar, 10 µm. C, Example eEPSCs at different holding potentials (C1) and I-V relationships (C2) of normalized AMPAR-mediated synaptic responses of D1+ or D1- MSN at different membrane holding potentials in D1-TdTomato or D1-ZsGreen rats after withdrawal from Sal or Coc SA [two-way ANOVA; Tukey’s multiple comparisons test: D1+ Sal vs. D1+ Coc at 20mV, 40mV, p<0.0001; D1+ Coc vs. D1- Sal at 20mV, 40mV, p<0.0001; D1+ Coc vs. D1- Coc at 40 mV, p=0.0114; * indicates significant main effects of cell-type/drug history and Vh, as well as Vh x cell-type/drug history interaction (all p<0.0001)]. D, Mean rectification index (RI) showing a higher RI in D1+ MSN from Coc rats compared to D1- MSN from Coc rats, D1+ MSN from Sal rats and D1- MSN from Sal rats [one-way ANOVA, Tukey’s multiple comparisons test: D1+ Sal vs. D1+ Coc, ****p<0.0001; D1+ Coc vs. D1- Sal, ****p<0.0001; D1+ Coc vs. D1- Coc, ****p<0.0001; D1+ Sal, 23 cells/9 rats (5M, 4F); D1+ Coc, 43 cells/12 rats (4M, 8F); D1- Sal, 10 cells/7 rats (4M, 3F); D1- Coc, 19 cells/11 rats (4M, 7F)]. E, Example eEPSCs (E1) and their time courses (E2) before and during 100 µM NASPM application in MSN from Coc or Sal rats [two-way ANOVA; Tukey’s multiple comparisons test, 15- 20 min for D1+ Sal vs. D1+ Coc, p<0.05; 15-20 min for D1+ Coc vs. D1- Coc, p<0.05; * indicates significant main effects of cell-type/drug history (p<0.0001) and time (p=0.0023), and a significant time x cell-type/drug history interaction (p<0.0001)]. F, Summary of mean eEPSC amplitudes after NASPM treatment showing increased NASPM sensitivity in D1+ MSN from Coc incubated rats [one-way ANOVA, Tukey’s multiple comparisons test: D1+ Sal vs. D1+ Sal, ***p=0.0003; D1+ Coc vs. D1- Coc, ***p=0.0002; D1+ Sal: 9 cells/5 rats (2M, 3F); D1+ Coc: 13 cells/7 rats (3M, 4F); D1- Coc: 6 cells/5 rats (3M, 2F)]. G-H, The I-V relationships of normalized eEPSCs at different membrane holding potentials and mean rectification index (RI) for each recorded MSN group. In Coc rats, DEAB-pretreated D1+ MSN showed a linear I-V relationship (two-way ANOVA; Tukey’s multiple comparisons test, at 20 mV and 40mV, D1+ CoC/Veh vs. D1+ Coc/DEAB, p<0.0001; D1+ Coc/Veh vs. D1- Coc/ DEAB, p<0.0001; D1+ Coc/Veh vs. D1- Coc/Veh, p<0.05). * indicates significant main effects of cell- type/treatment and Vh, as well as Vh x cell-type/treatment interaction (all p<0.0001) and a lower RI relative to Veh-pretreated D1+ MSN [one-way ANOVA, Tukey’s multiple comparisons test: D1+ Coc/Veh vs. D1+ Coc/DEAB, p<0.0001; D1+ Coc/Veh vs. D1- Coc/DEAB, p<0.0001; D1+ Coc/Veh vs. D1- Coc/Veh, p<0.0001; D1+ Coc/Veh, 15 cells/6 rats (4M, 2F); D1+ Coc/DEAB, 16 cells/6 rats (4M, 2F); D1- Coc/Veh, 7 cells/6 rats (3M, 3F); D1- Coc/DEAB, 11 cells/6 rats (4M, 2F)]. I-V relationship and RI of the DEAB-pretreated D1- MSN from Coc rats did not differ from Veh-pretreated MSN from Coc rats (p>0.05). I-K, D1+ MSN from drug-naïve D1-ZsGreen rats after RA pretreatment (2 µM for 1-2 h) showed inward rectification of I-V relationship [two-way ANOVA, * indicates significant main effects of cell- type and Vh, as well as Vh x cell-type interaction (all p<0.0001)], a higher RI [unpaired t-test, ****p<0.0001; D1+, 15 cells/5 rats (3M, 2F); D1-, 11 cells/5 rats (3M, 2F)] and increased NASPM sensitivity [unpaired t-test, *p=0.0196; D1+, 6 cell/4 rats (2M, 2F); D1-, 4 cells/4 rats (2M, 2F)] compared to D1- MSN.

These data indicate that incubation of cocaine craving is accompanied by CP-AMPAR upregulation selectively in D1 MSN. To determine if activation of RA signaling during incubation of cocaine craving also occurs only in D1 MSN, slices from D1-Cre/ZsGreen or TdTomato rats (WD40-50 after cocaine self-administration) were preincubated with the RA synthesis inhibitor DEAB (50 µM, 1-2 h) or its vehicle. We observed inward rectification, quantified as an elevated RI, in D1+ MSN in vehicle-pretreated slices from cocaine rats whereas no evidence of CP-AMPAR upregulation was observed in DEAB-pretreated slices in either D1+ or D1- MSN (Fig. 5G, two-way RM ANOVA, Vh x cell type/treatment interaction, F_(18,270)_=14.64, p<0.0001; main effect of Vh, F_(1.991, 89.58)_=20409, p<0.0001; main effect of cell type/treatment, F_(3, 45)_=16.72, p<0.0001; Fig. 5H, one-way ANOVA, F_(3,45)_=46.39, P<0.0001; Tukey’s multiple comparisons test, D1+ Veh vs. D1+ DEAB, p<0.0001; D1+ Veh vs. D1- DEAB, p<0.0001; D1+ Veh vs. D1- Veh, p<0.0001]. These results indicate that homeostatic plasticity leading to disinhibition of RA synthesis and CP-AMPAR upregulation occurs only in D1 MSN.

To determine if we could pharmacologically activate this pathway in both MSN subtypes, we prepared slices from drug-naïve D1-Cre/ZsGreen or TdTomato rats and pretreated the slices with RA (2 µM, 1-2 h) before recording AMPAR eEPSCs from D1+ and D1- MSN. We observed that RA pretreatment elicited inward rectification and increased NASPM sensitivity selectively in D1+ MSN (Fig. 5I, two-way ANOVA, Vh x cell type interaction, F_(6,126)_=15.54, p<0.0001; main effect of Vh, F_(6, 126)_=8064, p<0.0001; main effect of cell type, F_(1, 126)_=20.98, p<0.0001; Fig. 5J, unpaired t-test, t_(24)_=5.325, p<0.0001; Fig. 5K; D1+: 6 cells/4 rats (2M, 2F); D1-: 4 cells/4 rats (2M, 2F); Fig. 5K, unpaired t-test, t_(8)_=2.909, p=0.0196). Taken together, these results suggest that RA-dependent homeostatic plasticity occurs selectively in D1 MSN during incubation of cocaine craving (Fig. 6). Furthermore, even when exogenous RA is present, it does not mediate synaptic insertion of CP-AMPARs in D2 MSN.

## DISCUSSION

### RA-mediated homeostatic plasticity during cocaine withdrawal

The idea that homeostatic plasticity underlies strengthening of drug-related excitatory synaptic transmission during abstinence is not new (e.g., ^37^). It was originally based on evidence for high activity in glutamate pathways in the reward circuitry and particularly the NAc during drug use versus relatively lower activity in these pathways during abstinence (see below). This contrast would be expected to lead to a homeostatic strengthening of glutamate synapses in the NAc. This would be adaptive during abstinence when the level of glutamate release is low. But it becomes maladaptive when a cue is presented, because glutamate release elicited by the cue will now trigger a very strong synaptic and behavioral response (Fig. S5A).

CP-AMPARs are implicated in a number of mechanistically distinct types of homeostatic plasticity^38^. Here we identify a specific homeostatic plasticity mechanism, mediated by RA signaling^16^, that underlies CP-AMPAR-dependent strengthening of glutamate synapses in the NAcc during incubation of cocaine craving (see Fig. S5B for schematic). We found that increasing RA signaling in NAcc MSN from drug naïve rats rapidly upregulates CP-AMPARs, and that this pathway is operative only in D1 MSN. After incubation of cocaine craving, CP-AMPAR levels were elevated, consistent with our prior recordings from unidentified MSN (e.g., ^9, 13, 14, 34^), although the present results are the first to show that this occurs selectively in D1 MSN of NAcc (see below for discussion of NAc shell). Accordingly, the ability of exogenous RA to increase CP-AMPAR levels was occluded in D1 MSN from cocaine incubated animals. Instead, inhibition of RA synthesis reduced CP-AMPAR levels in these D1 MSN, suggesting that RA signaling is tonically active after cocaine withdrawal and that this required to maintain elevated synaptic levels of CP-AMPARs. Finally, recapitulating our in vitro results, we found that interruption of RA signaling in the NAcc of incubated rats reduces cue-induced cocaine seeking back to control levels. Our work adds to growing literature demonstrating a role for RA-mediated homeostatic plasticity in vivo ^39-43^.

### Expression mechanisms of incubation parallel those of RA-dependent homeostatic plasticity

While RA is an important transcriptional regulator ^17, 18^, RA-mediated homeostatic plasticity depends on translation not transcription, and specifically on increased GluA1 translation leading to synaptic insertion of homomeric GluA1 receptors ^24, 25, 28, 29, 44^. Our prior findings, using the same cocaine regimen used here, parallel this. First, we showed that CP-AMPARs upregulated after incubation are homomeric GluA1 ^9^. Then we showed that maintenance of elevated CP-AMPAR levels depends on ongoing protein translation, but not transcription ^45^. Finally, we directly measured translation in the NAcc using puromycin, which incorporates into newly translated proteins ^46^. GluA1 translation did not differ between control and cocaine rats on WD1, whereas it was elevated in cocaine rats after incubation. Cocaine withdrawal did not affect GluA2 translation or *Gria1* or *Gria2* mRNA levels measured in NAcc homogenates or synaptoneurosomes ^47^. Most recently, we demonstrated RA- and translation-dependent regulation of GluA1 surface expression in cultured NAc neurons ^48^. Together these data indicate that increased GluA1 translation, not transcription, contributes to incubation. The present results significantly extend these findings by demonstrating that manipulation of RA signaling regulates CP-AMPAR levels.

As noted above, CP-AMPAR upregulation and tonic RA signaling during incubation are confined to D1 MSN. Furthermore, in MSN from both drug-naïve and cocaine incubated rats, exogenous RA only influences CP-AMPAR levels in D1 MSN. These results suggest that an element(s) of the RA cascade is dysfunctional in D2 MSN or else counteracted by an opposing process. Future studies will pursue this. Selective upregulation of CP-AMPARs in D1 MSN is not unexpected given strong evidence that, although D1 and D2 MSN work in concert to govern motivated behavior, plasticity strengthening excitatory drive to D1 MSN promotes motivated behavior ^49-52^. As noted above, in prior studies of cocaine incubation we detected significant CP-AMPAR upregulation in NAcc even though we were recording unidentified MSN (e.g., ^9, 13, 14, 34^). One contributing factor may be that more striatal MSN express D1 than D2 receptors ^53, 54^.

### What events might activate RA signaling during incubation?

While our findings establish RA-mediated CP-AMPAR upregulation as the expression mechanism for incubation, the question remains of how RA signaling is activated during cocaine withdrawal. In culture models, RA-mediated homeostatic plasticity is triggered by a reduction in neuronal activation and Ca^2+^ signaling, leading to disinhibition of RA synthesis ^16^. We propose that the cumulative effect of multiple events occurring in NAcc over the first weeks of cocaine withdrawal ultimately lowers Ca^2+^ activity in MSN below the threshold for disinhibition of RA synthesis. Supporting this idea, several lines of evidence indicate decreased activity in brain areas sending excitatory inputs to the NAc (and in the NAc itself) after discontinuing cocaine self-administration ^55-63^. The only one of these studies that specifically studied incubation of cocaine craving found a negative correlation between resting local field potential power in the NAcc (gamma frequency) measured on WD1 and the magnitude of incubation on WD30 ^60^, suggesting that a reduction of NAc activity in very early withdrawal is related to incubation.

While consistent with a homeostatic plasticity hypothesis, the observation of reduced NAcc activity on WD1 begs the question of why weeks of withdrawal are required to observe CP-AMPAR elevation. We propose that additional events are required before Ca^2+^ activity in MSN is sufficiently reduced to trigger RA synthesis. There are two strong candidates, both detected after the same cocaine regimen used here. First, after 2-3 weeks of withdrawal, we have observed synaptic insertion of atypical NMDARs containing the GluN3 subunit and shown that this is necessary for subsequent incubation ^64^. The presence of GluN3 in triheteromeric NMDARs endows them with atypical properties including low Ca^2+^ permeability ^65-68^. Thus, insertion of GluN3-NMDARs likely explains our observation of reduced NMDAR-mediated Ca^2+^ entry into NAcc MSN dendritic spines after incubation of cocaine craving ^64, 69^. Second, between WD14 and WD25, we found that surface expression of mGlu1, which is canonically linked to Ca^2+^ mobilization ^70^, decreases just before CP-AMPARs increase and that this decrease is essential for CP-AMPAR accumulation and incubation ^13^. Overall, these results support the idea that multiple “hits” over the first weeks of cocaine withdrawal reduce MSN activity to a threshold that ultimately activates RA signaling.

In the NAc shell, a cocaine-induced reduction in membrane excitability due to changes in several conductances is well documented ^71-73^. This occurs very early in withdrawal from regimens leading to incubation of cocaine craving in rats ^74^ and, along with silent-synapse based remodeling ^75^, contributes to a synapse-membrane homeostatic cascade in NAc shell that is critical for incubation and culminates in pathway-specific CP-AMPAR upregulation ^74^. It is not known if the excitability of NAcc MSN in rats is reduced during incubation, but in mice a cocaine self-administration regimen that reduces MSN excitability in the shell on WD1 does not alter excitability in the core ^76^. Other evidence also suggests distinct incubation-related adaptations in the two subregions, despite the commonality of CP-AMPAR upregulation ^64, 74^. Thus, the involvement of RA-mediated homeostatic plasticity may be unique to NAcc. It is also interesting to note that, although recordings from unidentified NAc shell MSN indicate low CP-AMPAR levels under baseline conditions, optogenetic studies have found significant levels in certain pathways^77, 78^. Furthermore, while the initial studies demonstrating incubation-related CP-AMPAR upregulation in NAc shell did not distinguish D1 and D2 MSN^79, 80^, subsequent work indicates that D1 vs D2 involvement depends on the cocaine regimen and the pathway ^81, 82^.

While the present study focused on manipulating RA after incubation had plateaued in order to test its role in maintaining CP-AMPARs and incubation, other recent studies are testing the effect of manipulating RA during the rising phase of incubation. Results indicate that enhancing RA signaling in the NAcc by administering shRNA against the RA degradative enzyme Cyp26b1 accelerated CP-AMPAR accumulation and the incubation of cocaine craving. Conversely, inhibiting RA synthesis via shRNA against the RA synthetic enzyme Aldh1a1 normalized CP-AMPAR levels in infected MSN but did not prevent incubation; the latter may reflect the ability of non-infected cells to support incubation ^83^.

### Role for RA regulation of transcription in addiction models

Although our work focuses on a non-transcriptional role of RA, there is also evidence that RA-mediated regulation of transcription contributes to addiction. In a study on the interaction between cocaine self-administration and environmental enrichment, transcriptomics and pathway analysis suggested a prominent role for RA signaling; furthermore, cocaine seeking in early withdrawal was augmented after knocking down the RA degradative enzyme Cyp26b1 in the NAc shell ^19^. A follow-up study showed that knockdown in the male rat NAc shell of fatty acid binding protein 5 (FABP5), which helps translocate RA from cytoplasm to the nucleus, decreased cocaine self-administration and altered MSN intrinsic excitability (but did not affect excitatory synaptic transmission) ^20^. It should be noted that this transcript is enriched in shell with much lower expression in the core subregion ^20^, and due to its role in translocation to the nucleus it is unlikely to contribute to dendritic RA signaling. Another study on cocaine seeking during withdrawal from environmental enrichment vs isolation found trends for a relationship between incubation (seeking in isolated rats on WD1 vs WD21) and RA-related pathways (‘RAR Activation’ and ‘PPARα/RXRα Activation; RXRs heterodimerize with RARs to regulate transcription), along with significant effects on these pathways for other contrasts ^84^, while a follow-up study of differentially expressed microRNAs between high and low cocaine-seeking isolated rats on WD21 found that 15 of these microRNAs targeted retinoic acid receptor-related orphan receptor B ^85^.

In NAcc, transcriptomic studies related to addiction have implicated RXRs and particularly RXRα. While RXRα transcript levels were not altered in the NAcc by cocaine self-administration, they were positively correlated with an “addiction index” derived from elements of cocaine self-administration behavior ^21^. A follow-up study found that RXRα controls transcriptional programs in both D1 and D2 NAcc MSN that regulate their excitability and that manipulating RXRα expression influenced cocaine CPP and self-administration of low cocaine doses; furthermore, systemic treatment with the RXR inhibitor HX531 reduced behavioral responses to cocaine (CPP and lever pressing) ^22^. There is no evidence for RXR function outside the nucleus, so our observations cannot be attributed to RXR. However, it will be exciting to explore how nuclear RAR and RXR transcriptional regulation may interact with dendritic RAR regulation to control motivated behavior.

### Broader potential effects of RA in NAcc dendrites

Paralleling what was found in hippocampal slices ^24^, our results indicate that RA-mediated homeostatic plasticity of excitatory transmission is associated with postsynaptic strengthening in the absence of evidence for changes in presynaptic function. However, there are many other aspects of synaptic transmission in the NAcc that should be explored in the future. In other brain regions, RA-mediated homeostatic plasticity not only enhances excitatory transmission but also oppositely regulates inhibitory synaptic transmission; in both cases, this is a non-transcriptional effect that requires protein translation ^24, 39, 86^. By regulating both excitatory and inhibitory transmission, RA alters the E/I balance and may thereby elicit metaplasticity that influences subsequent Hebbian plasticity ^87^. Future studies should explore the potential role of these mechanisms in incubation of cocaine craving.

## Acknowledgements

This work was supported by USPHS grants R01 DA049930 (M.E.W.) and F32 DA046141 (A.M.W.). We thank Alana L. Moutier, Jonathan R. Funke and Madelyn M. Beutler for assistance with drug self-administration and Dr. Daniel T. Christian for assistance with pilot experiments.

## Conflict of interest statement

Dr. Wolf and OHSU have a financial interest in Eleutheria Pharmaceuticals LLC, a company that may have a commercial interest in results related to the research described herein. This potential conflict of interest has been reviewed and managed by OHSU. The other authors declare no competing interests.

**Supplementary information is available at MP’s website.**

