## Supplemental info for "Retinoic acid-mediated homeostatic plasticity drives cell type-specific CP-AMPAR accumulation in nucleus accumbens core and incubation of cocaine craving"

**Contents**: Figures S1-S5; Supplementary Materials and Methods; References for Supplementary Materials and Methods

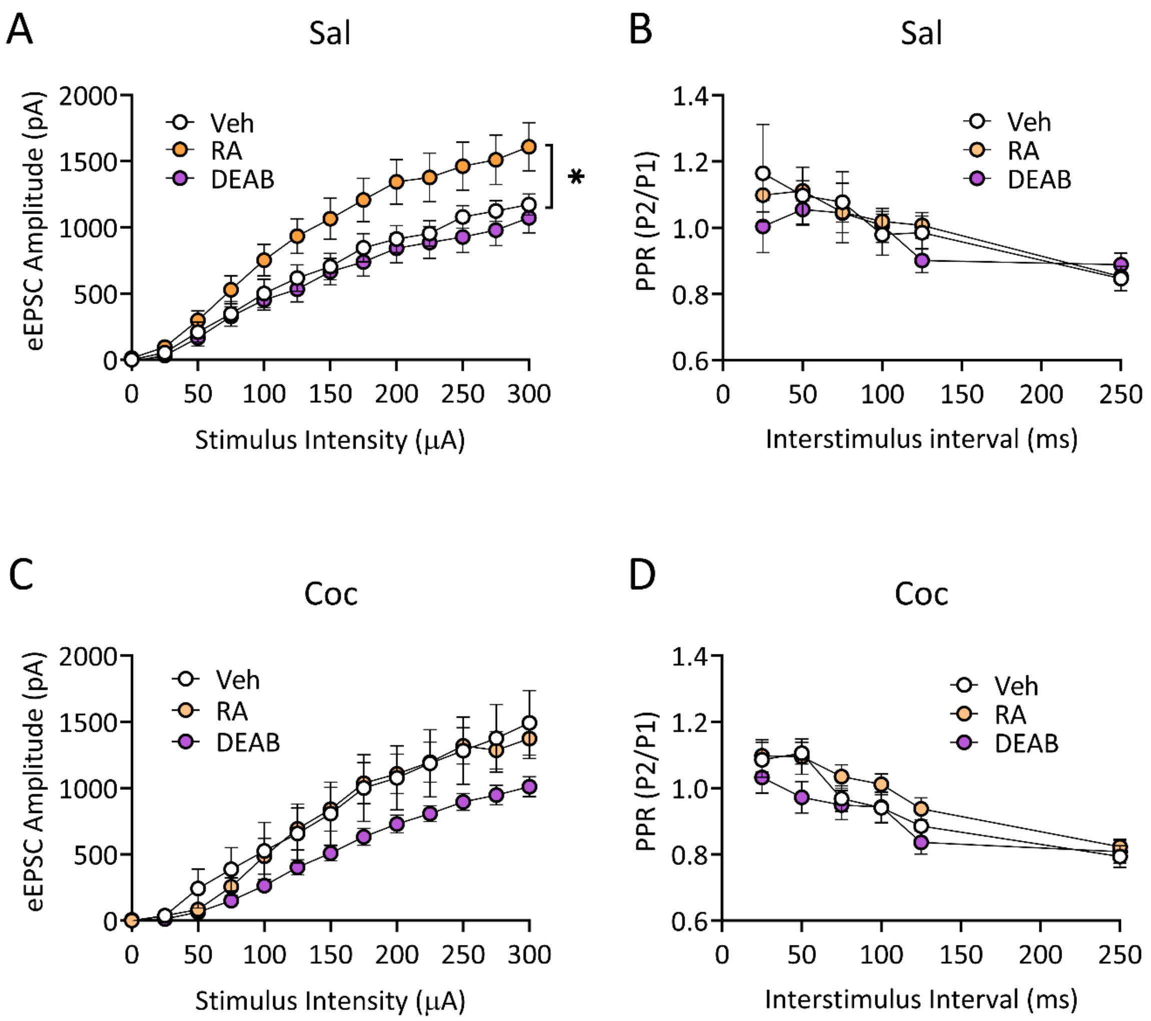


**Fig. S1. Retinoic acid (RA)-mediated increase in excitatory synaptic transmission occurs without presynaptic changes in nucleus accumbens (NAc) core medium spiny neurons (MSN) from saline (Sal) or cocaine (Coc) wild type rats**

A, I-O plots of eEPSCs recorded with incremental electrical stimulation (0-300 μA) show increased synaptic strength after RA pretreatment in MSN from Sal rats compared to Veh or DEAB pretreatment [two-way ANOVA, stimulus intensity x treatment interaction, $F_{(24,312)}=3.004$, $p<0.0001$; main effect of treatment, $F_{(2,26)}=3.929$, *$p=0.0323$; Tukey's multiple comparisons test, Veh vs. RA: 150-300 μA, $p<0.05$; RA vs. DEAB: 125-150 μA, $p<0.05$, 175-300 μA, $p<0.01$; Veh, 10 cells/4 rats (2M, 2F); RA, 9 cells/4 rats (2M, 2F); DEAB, 10 cells/4 rats (2M, 2F)].

B, Paired-pulse ratios did not differ among treatment groups [two-way ANOVA, ISI x treatment interaction, $F_{(10,95)}=1.261$, $p=0.2639$; main effect of treatment, $F_{(2,19)}=0.1826$, $p=0.8345$; Veh, 7 cells/4 rats (2M, 2F); RA, 9 cells/4 rats (2M, 2F); DEAB, 6 cells/3 rats (2M, 1F)).

C, I-O plots of eEPSCs recorded with incremental electrical stimulation (0-300 μA) show increased synaptic strength after RA pretreatment in MSN from Coc rats [two-way ANOVA, stimulus intensity x treatment interaction, $F_{(24,420)}=2.262$, $p=0.0007$; main effect of treatment, $F_{(2,35)}=2.040$, $p=0.1451$; Tukey's multiple comparisons test: Veh vs. DEAB, 250-300 μA, $p<0.05$; RA vs DEAB, 250 μA, $p<0.05$; Veh, 11 cells/5 rats (3M, 2F); RA, 11 cells/6 rats (3M, 3F); DEAB, 16 cells/7 rats (3M, 4F)].

D, Paired-pulse ratios did not differ among treatment groups [two-way ANOVA, ISI x treatment interaction, $F_{(10,185)}=1.175$, $p=0.3100$; main effect of treatment, $F_{(2,37)}=1.440$, $p=0.2499$; Veh, 11 cells/5 rats (3M, 2F); RA, 13 cells/6 rats (3M, 3F); DEAB, 16 cells/7 rats (3M, 4F)].

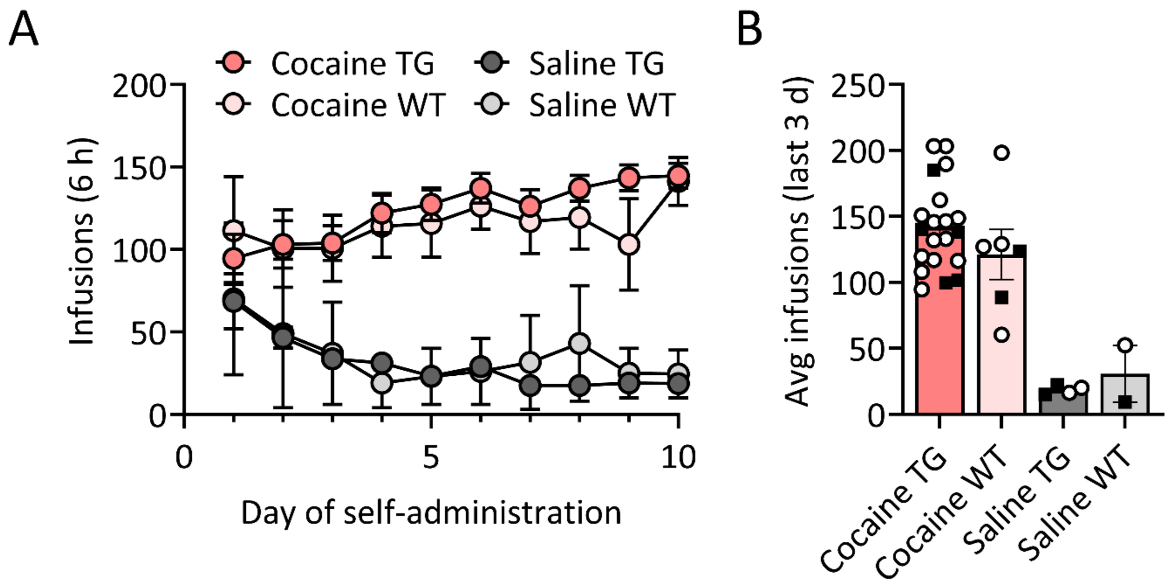


**Fig. S2. Comparison of cocaine or saline infusions between wild-type (WT) and transgenic rats (TG) over the 10 days self-administration (Fig. 3D reanalyzed to disaggregate WT and TG data).**

A, The number of cocaine or saline infusions taken over the 10 days of self-administration did not differ between WT and TG rats [two-way ANOVA, day x drug interaction, $F_{(27,252)}$=1.753, p=0.0144; main effect of day, $F_{(2.204,61.71)}$=0.4935, p=0.6310; main effect of drug, $F_{(3,28)}$=11.72, p<0.0001; Sidak's multiple comparisons post hoc test: Coc TG vs Coc WT, p>0.05; Sal TG vs. Sal WT, p>0.05; Cocaine WT, 6 rats (2M, 4F), TG, 20 rats (5M, 15F); Saline, 2 rats (1M, 1F); TG, 9 rats (5M, 4F)].

B, The average number of cocaine or saline infusions in the last 3 days of cocaine self-administration did not differ between WT and TG rats [two-way ANOVA, rat type x treatment interaction, $F_{(1,28)}$=0.9623, p=0.3350; main effect of rat type, $F_{(1,28)}$=0.05894, p=0.8100; main effect of treatment, $F_{(1,28)}$=40.34, p<0.0001; Sidak's multiple comparisons post hoc test: Coc TG vs Coc WT, p>0.05; Sal TG vs. Sal WT, p>0.05]. Data presented as mean ± S.E.M. Individual data points for each rat (■ Male, O Female) are included in bar graphs.

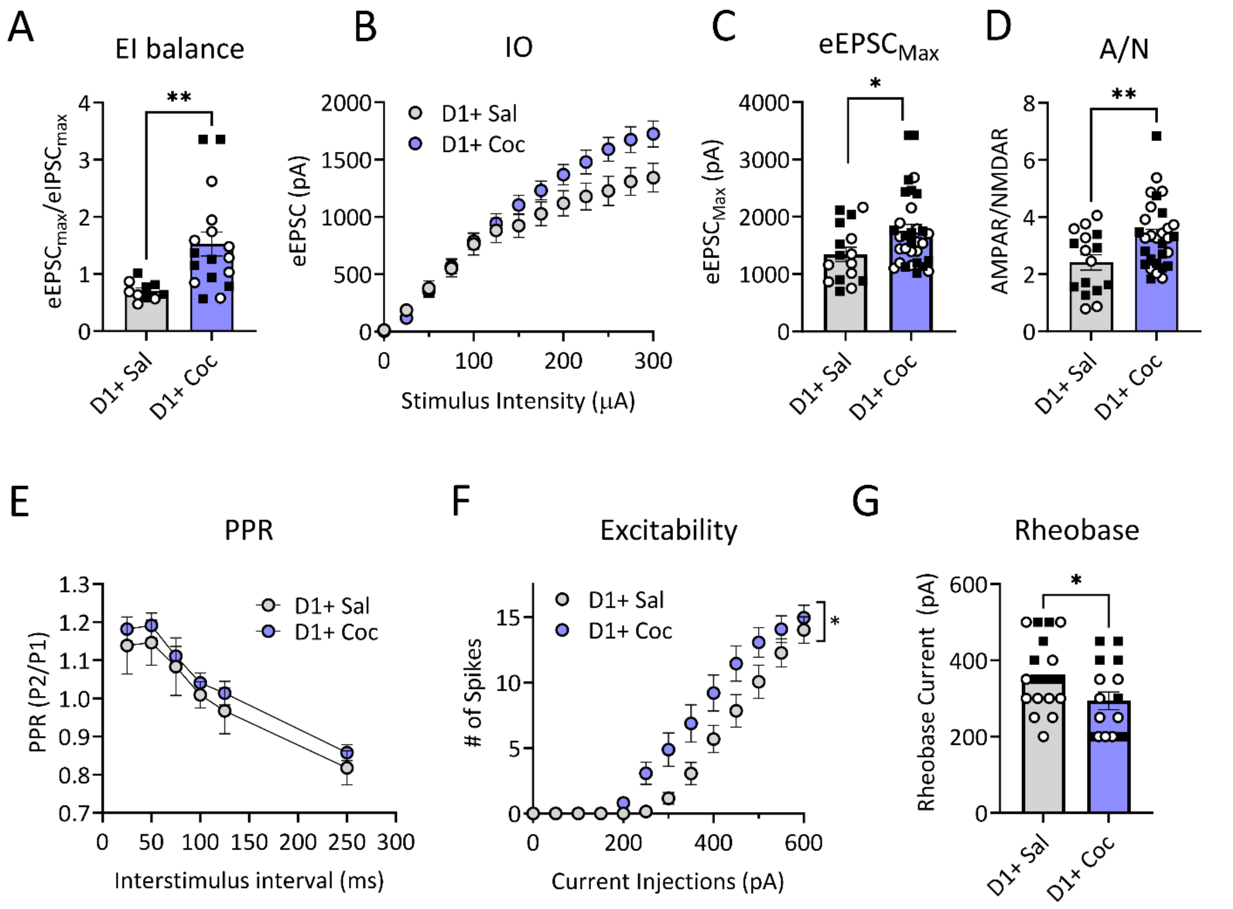


**Fig. S3. Increased excitatory tone in nucleus accumbens (NAc) core D1 medium spiny neurons (MSN) after >40 days of withdrawal from cocaine self-administration.**

A, E-I ratios (averages of eEPSCs recorded at -70 mV/eIPSCs recorded at 0 mV in the same cell using the same electrical intensity) for each D1+ MSN from saline (Sal) and cocaine (Coc) rats, showing increased EI balance in D1 MSN from Coc rats compared to D1 MSN from Sal rats [unpaired t-test, $t_{(25)}$=2.949,**p=0.0068; D1+ Sal, 10 cells/4 rats (2M, 2F); D1+ Coc, 17 cells/6 rats (3M, 3F)].

B, I-O plots of eEPSCs recorded with incremental electrical stimulation (0-300 μA) [D1+ Sal, 16 cells/6 rats (3M, 3F); D1+ Coc, 34 cells/8 rats (4M, 4F)].

C, Comparison of maximal eEPSC amplitudes evoked by the 300 μA stimulation in D1 MSN from Sal and Coc rats [unpaired t-test, $t_{(46)}$=2.219, *p=0.0314; D1+ Sal, 16 cells/6 rats (3M, 3F); D1+ Coc, 32 cells/8 rats (4M, 4F)].

D, AMPAR/NMDAR ratios for each group. D1 MSN from Coc rats showed a higher A/N ratio than D1 MSN from Sal rats [unpaired t-test, $t_{(44)}$=2.700, **p=0.0098; D1+ Sal, 16 cells/6 rats (3M, 3F); D1+ Coc, 30 cells/8 rats (4M, 4F)].

E, No difference in paired-pulse ratios of eEPSCs at different inter stimulus intervals (ISIs) for each group [D1+ Sal, 7 cells/4 rats (2M, 2F); D1+ Coc, 21 cells/7 rats (3M, 4F)].

F, Summary of D1 MSN neuronal excitability for Sal vs Coc rats [two-way RM ANOVA, injected current x treatment interaction, $F_{(12,396)}$=3.489, p<0.0001; main effect of injected current, $F_{(1.653, 54.55)}$=162.4, p<0.0001; main effect of treatment, $F_{(1, 33)}$=4.749, p=0.0366; D1+ Sal, 19 cells/5 rats (2M, 3F); D1+ Coc, 16 cells/6 rats (3M, 3F)].

G, Comparison of mean rheobase currents in D1 MSN from Sal and Coc rats [unpaired t-test, $t_{(33)}$=2.187, *p=0.0359; D1+ Sal, 19 cells/5 rats (2M, 3F); D1+ Coc, 16 cells/6 rats (3M, 3F)].

Data are presented as mean ± S.E.M. Individual data points for each rat are included in bar graphs (■ Male, O Female).

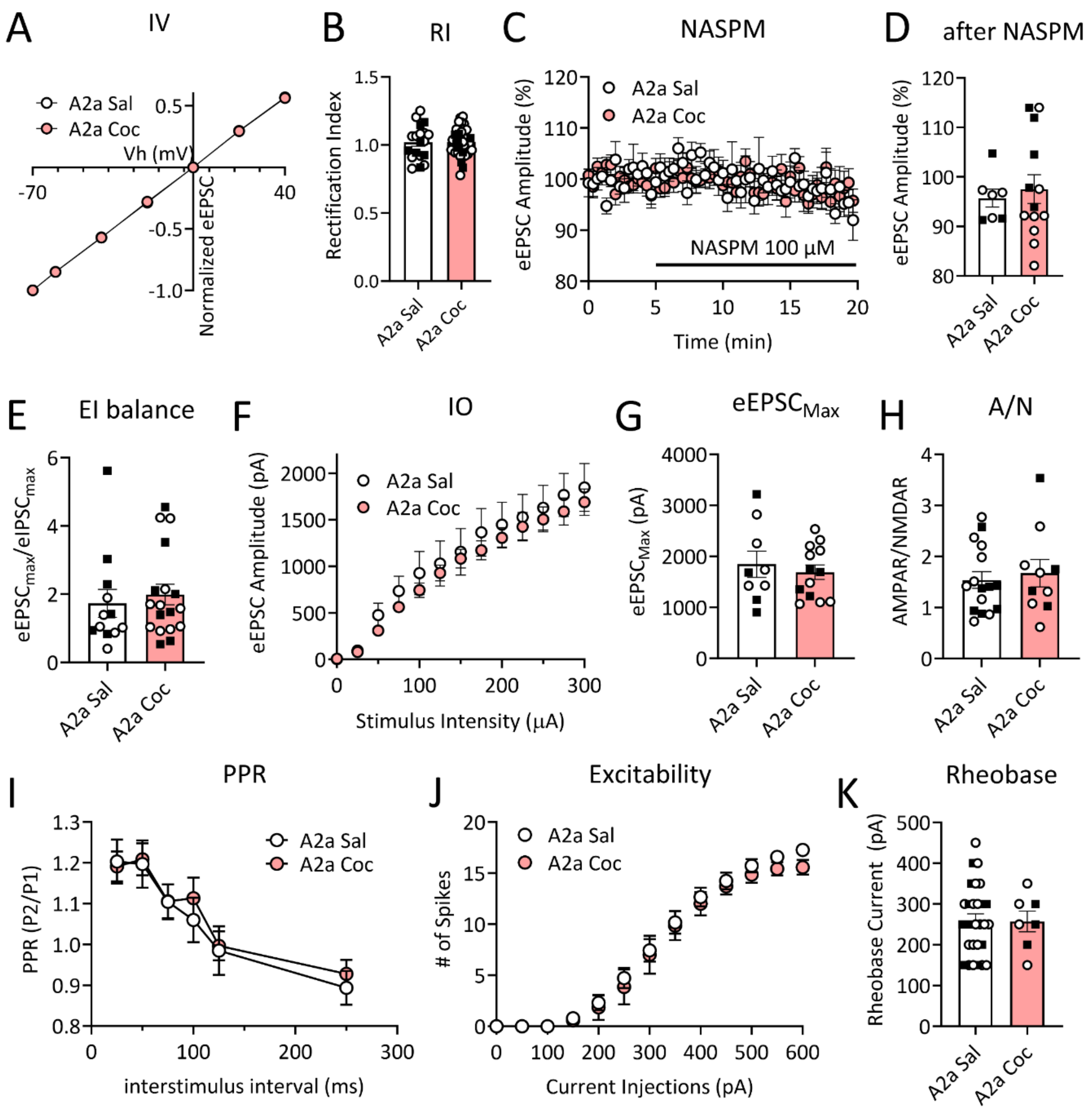


**Fig. S4. No alteration in neuronal or synaptic properties of A2a+ medium spiny neurons (MSN) in nucleus accumbens core (NAcc) after >40 days of withdrawal from cocaine (Coc) self-administration (SA).**

A, The I-V relationships of normalized AMPAR-mediated synaptic responses of A2a+ MSN at different membrane holding potentials after withdrawal from saline (Sal) or Coc SA (two-way ANOVA, no significant drug x Vh interaction or main effect of drug, p>0.05). A2a+ MSN were identified by crossing A2a-Cre rats with TdTomato or ZsGreen reporter lines.

B, Mean rectification index (RI) did not differ between A2a+ MSN from Sal and Coc rats [unpaired t-test, $p>0.05$; Sal, 19 cells/6 rats (2M, 4F); Coc, 36 cells/6 rats (2M, 4F)].
C, Time course of eEPSC amplitude before and during 100 μM NASPM application in A2a+ MSN from Sal or Coc rats (two-way ANOVA, all $p>0.05$).
D, Summary of NASPM effects on eEPSC amplitude in each group, measured after 15-20 min of NASPM application, showing no difference in NASPM sensitivity between A2a+ MSN from Sal or Coc rats [unpaired t-test, $p>0.05$; Sal, 7 cells/5 rats (2M, 3F); Coc, 13 cells/7 rats (3M, 4F)].
E, E-I ratios (averages of eEPSCs recorded at -70 mV/eIPSCs recorded at 0 mV in the same cell using the same electrical intensity, 300 μA) for each A2a+ MSN from Sal and Coc rats, showing no difference in EI balance between A2a+ MSN from Sal rats and Coc rats [unpaired t-test, $p>0.05$: Sal, 12 cells/5 rats (3M, 2F); Coc, 18 cells/5 rats (2M, 3F)].
F, Mean input-output (IO) relationship of AMPAR-mediated eEPSCs in response to incremental electrical stimulation showing no difference in the strength of synaptic responses between A2a+ MSN from Sal and Coc rats [Sal, 9 cells/5 rats (2M, 3F); Coc, 13 cells/6 rats (2M, 4F)].
G, Comparisons of maximal eEPSC amplitudes evoked by the 300 μA stimulation in A2a+ MSN from Sal and Coc rats (unpaired t-test, $p>0.05$).
H, AMPAR/NMDAR ratios for each group. No difference was observed between groups [Sal, 16 cells/5 rats (2M, 3F); Coc, 10 cells/5 rats (2M, 3F)].
I, Paired-pulse ratios (PPR) of eEPSCs at different inter stimulus intervals (ISIs) in NAcc A2a+ MSN do not differ between Sal and Coc rats [two-way ANOVA, all $p>0.05$; Sal, 11 cells/5 rats (3M, 2F); Coc, 10 cells/4 rats (2M, 2F)].
J, Mean number of action potentials in response to incremental depolarizing current injections (0 pA to 600 pA/Δ50 pA, 0.5 s) in A2a+ MSN [two-way ANOVA, all $p>0.05$; Sal, 29 cells/6 rats (2M, 4F); Coc, 7 cells/4 rats (2M, 2F)].
K, The rheobase currents measured using step current injections were similar in each group of A2a+ MSN [unpaired t-test, $p>0.05$; Sal, 29 cells/6 rats (2M, 4F); Coc, 7 cells/4 rats (2M, 2F)].
Data are presented as mean ± S.E.M. Individual data points for each rat are included in bar graphs (■ Male, O Female).

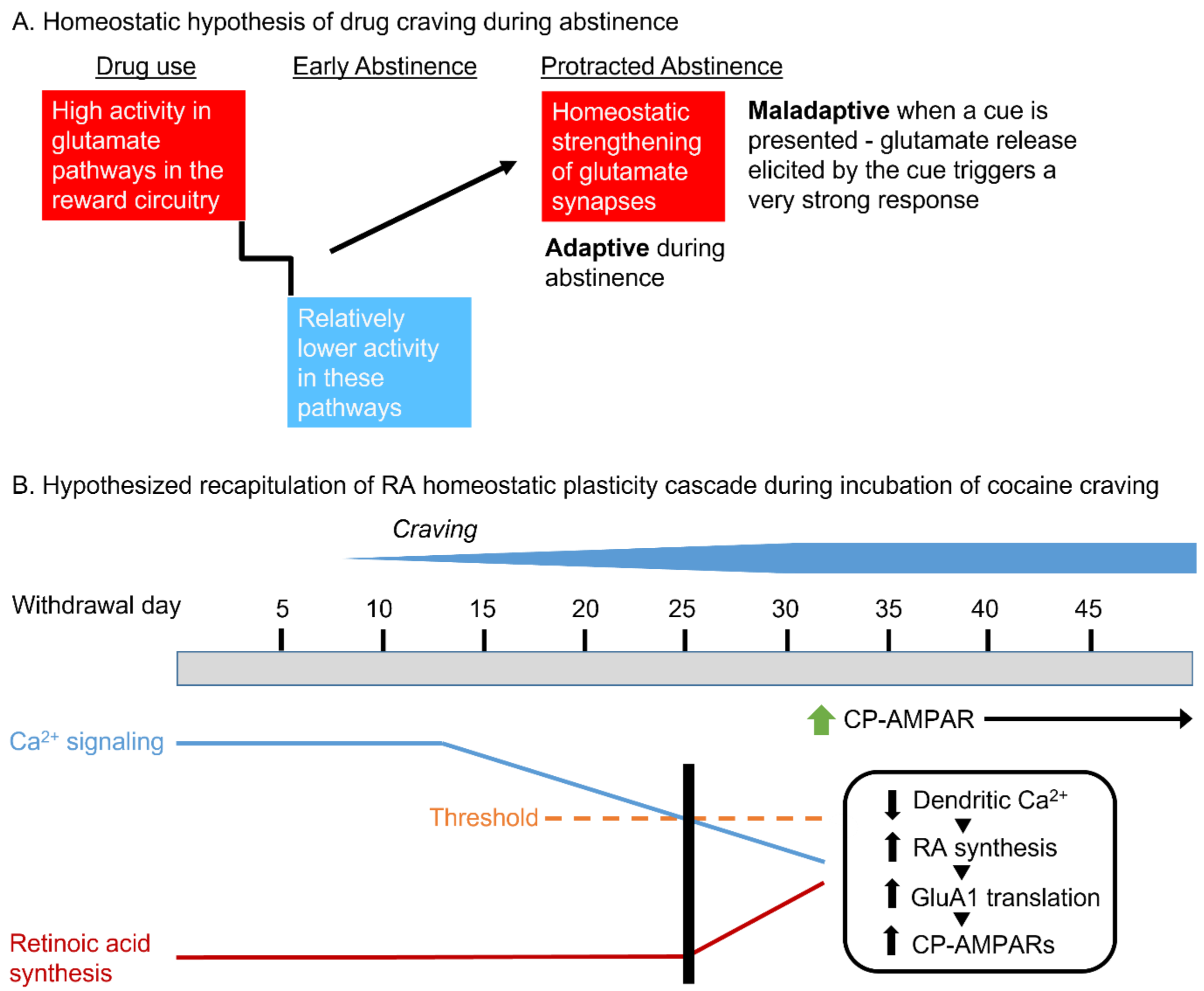


**Fig. S5. Proposed mechanism through which retinoic acid (RA)-dependent homeostatic strengthening of glutamatergic synapses occurs in D1 medium spiny neurons (MSN) during incubation of cocaine craving.**

A. Adaptations strengthening glutamate synapses in NAc core (NAcc) MSN during abstinence from cocaine self-administration are hypothesized to result from homeostatic plasticity.

B. The cumulative effect of multiple events occurring in NAcc over the first weeks of forced abstinence from cocaine self-administration is proposed to lower $Ca^{2+}$ activity in MSN below the threshold for disinhibition of RA synthesis. These events include decreased activity in brain areas sending excitatory inputs to the NAcc (and in the NAcc itself), reduced mGlu1 tone, and synaptic incorporation of GluN3-NMDARs (see Discussion for details). Increased RA signaling in NAcc MSN during forced abstinence leads to increased GluA1 translation, thereby contributing to the increase in synaptic CP-AMPAR levels that mediates incubated cocaine seeking. This cascade preferentially occurs in NAcc D1 MSN.

**Supplementary Materials and Methods**

Subjects. All procedures were approved by the Oregon Health & Science University Institutional Animal Care and Use Committee in accordance with the USPHS Guide for Care and Use of Laboratory Animals. We used wildtype (WT) and transgenic Long-Evans rats generated in our breeding colony; breeder rats for transgenic lines are obtained from the Rat Resource & Research Center (RRRC) and wildtype breeders from Charles River (Crl:LE, strain 006). Some experimental WT rats were obtained directly from Charles River. To enable selective manipulation of D1 and D2 receptor expressing MSN in the NAcc, we used knock-in rat lines, generated by CRISPR-Cas9 gene editing, encoding iCre recombinase immediately after the Drd1a or adenosine 2a receptor (Adora2a; A2a) loci.  A2a was targeted because it is selectively expressed in D2 MSN, while the D2 receptor is also expressed on cholinergic interneurons and other elements [1]. These transgenic rat lines [LE-Drd1$^{\text{em1(iCre)Berke}}$ (RRRC #856) and LE-Adora2a$^{\text{em1(iCre)Berke}}$ (RRRC #857)], hereafter termed D1-Cre and A2a-Cre, have been thoroughly validated. Whole genome sequencing, FISH, and Cre-driven viral expression confirmed correctly targeted Cre expression, while behavioral studies confirmed normal learning, motivation, and locomotor responses to cocaine [2]. In order to identify Cre+ neurons for slice recordings, D1-Cre or A2a-Cre rats were bred to reporter lines expressing either ZsGreen [Tg$^{(\text{CAG-}\textit{lox}\text{P-STOP-}\textit{lox}\text{P-ZsGreen)561Bryd}}$, RRRC #797] or TdTomato [LE-Rosa26$^{\text{(CAG-LSL-TdTomato)em1Rrrc}}$, RRRC #938] in the presence of Cre. Rats were allowed free access to food and water and maintained on a 12-h light/dark cycle. Rats were 10-15 weeks old at the onset of the experiment.

*Surgery.* Rats destined for drug self-administration were anesthetized with isoflurane (MWI Animal Health, Boise, ID) for surgery to implant a jugular catheter. The catheter consists of a mesh-reinforced, 22-gauge cannula (placed on the back, ~1 cm caudal to the shoulder blades; Plastics One, Roanoke, VA) which has an attached 10 cm silastic tube (The Dow Chemical Company, Midland, MI). The tube is threaded subcutaneously around the shoulder blade and inserted into the jugular vein. Meloxicam (5 mg/kg, s.c.; Covetrus, Portland, ME) was administered as an analgesic pre- and post-operatively. Following catheter surgery, rats were single-housed with enrichment (nylabones and nesting packs) for the duration of the experiment. During recovery from surgery (5-7 days) and during the 10 days of cocaine self-administration, catheters were flushed every 24-48 h with 0.9% sterile saline (Baxter International, Deerfield, IL) containing the β-lactam antibiotic Cefazolin (0.1 ml of 100 mg/mL, IV; Covetrus). Some rats, destined for intra-NAcc infusion of vehicle or DEAB prior to cue-induced seeking tests, also underwent stereotaxic surgery to implant bilateral intracranial guide cannulas aimed at the NAcc (+1.4 A/P, +2.35 M/L, -6.35 D/V) as described previously [3, 4].

*Cocaine self-administration.* Rats self-administered cocaine (0.5 mg/kg/infusion; obtained from the NIDA Drug Supply Program) or 0.9% saline (control condition) for 6 h/d for 10 sessions across 12 d (2 days off following the 5$^{th}$ session) under a fixed ratio 1 schedule as described previously (e.g., [5-8]). Self-administration sessions were carried out in operant chambers with two nose-poke holes located on opposite walls. Active hole pokes triggered the infusion pump and simultaneously activated a 4 s cue light (simultaneous 4 s timeout period). Inactive pokes had no consequence. Following self-administration training, rats were singly housed and handled once per week for the duration of forced abstinence.

*Cue-induced seeking tests.* Some rats, implanted with both jugular catheters and bilateral intracranial guide cannulas aimed at NAcc (see Surgery), received cue-induced seeking tests on withdrawal day (WD) 1 and WD50-60 following extended-access cocaine self-administration. The WD1 test was performed without manipulation to establish baseline craving. Rats were placed in the self-administration chamber and allowed to freely poke in either the previously active or inactive hole for 1 h under extinction conditions [i.e., a poke in the active hole resulted in presentation of the same 4 s cue light previously paired with cocaine (or saline) but the rat received no cocaine and a poke in the inactive hole continued to have no consequence]. The number of pokes in the previously active hole was used as a measure of cocaine seeking or craving. Beam breaks recorded by near infrared sensors in the operant boxes provided a measure of locomotor activity during seeking tests. Following the WD1 seeking test, rats were returned to home cages. Rats habituated to injection procedure 4 days prior to intra-NAcc infusions. One hour prior to the WD50-60 seeking test, the same rats received bilateral infusion (0.5 µl/side) of vehicle (0.1% DMSO in aCSF) or DEAB (50 µM in 0.1% DMSO + aCSF) into the NAcc. As described previously [3], a syringe pump was connected to a 10-µl Hamilton syringe, which was then connected to 30-ga injectors via PE20 tubing (Fisher Scientific; Hampton, NH). Tips of injectors extended 1 mm below the tip of the guide cannula and solutions were infused at a rate of 0.5 µl/min. Injectors were left in place for an additional minute to allow for diffusion away from the injection site. One hour after the injection, rats were placed in the self-administration chamber and the cue-induced seeking test was performed as described for WD1.

*Slice electrophysiology.* Electrophysiological experiments were performed as we have described previously (e.g., [5, 6, 8-10]). Cocaine and saline rats were recorded between WD40 and WD60. Briefly, rats were anesthetized with isoflurane and brains were rapidly removed. Coronal slices at the level of the NAc (280 μm) were cut with a vibrating microtome in ice-cold N-methyl-D-glucamine (NMDG)-based cutting solution (in mM: 93 NMDG, 2.5 KCl, 1.2 $NaH_2PO4$, 30 $NaHCO_3$, 20 HEPES, 25 glucose, 5 sodium ascorbate, 3 sodium pyruvate, 10 $MgCl_2$, 0.5 $CaCl_2$) and then incubated at 32-34°C for 15 min. Slices were then transferred to a holding chamber that contained room temperature, oxygenated (95% $O_2$/5% $CO_2$) artificial cerebral spinal fluid [aCSF; in mM: 109 NaCl, 4.5KCl, 1 $MgCl_2$, 2.5 $CaCl_2$, 1.2 $NaH_2PO_4$, 35 $NaHCO_3$, 11 glucose, 20 HEPES, 0.4 sodium ascorbate (pH7.3, 300-310 mOsm)] until use. Whole cell patch clamp recordings of MSN were conducted at least 1 h after slicing in oxygenated (95% O2/5% CO2) and warm (30-32°C) aCSF [in mM: 126 NaCl, 3 KCl, 1.5 $MgCl_2$, 2.4 $CaCl_2$, 1.2 $NaH_2PO_4$, 11 glucose, 26 $NaHCO_3$]. Picrotoxin (0.1 mM) and APV ((2R)-amino-5-phosphonopentanoate) (0.05 mM) were added into the recording aCSF to pharmacologically isolate AMPAR transmission. Recordings were conducted using patch pipettes (2–3 MΩ) filled with a Cs-based/spermine-containing internal solution (in mM: 115 $CsMeSO_3$, 10 HEPES, 20 CsCl, 0.6 EGTA, 7.5 QX-314, 10 NaPhosphocreatine, 4 MgATP, 0.4 NaGTP, 0.1 spermine; adjusted to pH7.3 and 290 mOsm). The liquid junction potential (~13 mV) for $CsMeSO_3$ recordings was corrected for current-voltage relationship measurements. The reversal potentials were stable (±2 mV) on different days and corrected to 0 mV. Spermine (100 μM) was added fresh to the internal solution each day. A bipolar tungsten stimulating electrode (FHC, Bowdoin, ME) placed ~200 μm from the recording site was used to elicit excitatory postsynaptic currents (EPSC) in MSN. Only neurons that exhibited stable baseline synaptic responses (<15% variability, 15 min) were included. The rectification index (RI), calculated as $[EPSC_{-70\ mV}/(-70 - E_{rev})]/[EPSC_{+40\ mV}/(+40-E_{rev})]$ [11], was used to assess the contribution of CP-AMPAR to synaptic transmission. For current clamp recordings, recording pipettes were filled with K-gluconate based internal solution (in mM): 140 K-gluconate, 10 HEPES, 0.2 EGTA, 2 $MgCl_2$, 0.1 $CaCl_2$, 4 $Na_2$-ATP, 0.3 Na-GTP; adjusted to pH7.3 and 290 mOsm. The intrinsic excitability evaluations were performed at -80 mV by adjusting membrane potential (typically less than ±50 pA current injection) in current clamp mode. For analysis of excitability measurements, each protocol was repeated three times and averaged.

*Reagents.* The following reagents to manipulate RA signaling were obtained from Sigma–Aldrich (St. Louis, MO): RA (R2625), DEAB (D86256), AM580 (A8843), and Ro 41-5253 (SML0573). DL-AP5 (HB0252), Picrotoxin (HB0506) and NASPM (HB0441) were obtained from Hello Bio (Princeton, NJ).

*Statistical Analyses.* Clampfit (v11, Molecular Devices) and GraphPad Prism10 were used for figures and statistical analyses. All data were assessed for normality using Shapiro-Wilk tests. For comparison of two groups, Student's t-tests (independent unless otherwise indicated) were used for normally distributed data and Mann Whitney test was used for nonnormally distributed data sets. Either one-way or two-way ANOVAs were utilized for comparing multiple groups followed by Tukey's or Bonferroni post hoc multiple comparisons. IV curves were analyzed using two-way repeated measures ANOVA with treatment as between-subject factor and voltage as within-subject factor. Rectification indexes were analyzed using unpaired t-tests for two group comparison or one-way ANOVA with Tukey's or Bonferroni post hoc multiple comparisons to compare more than two groups. The number active pokes, inactive pokes and infusions during self-administration and cue-induced seeking tests were analyzed with two-way Mixed ANOVA with day of SA or WD day as within-subject factors and drug history as between-subject factor, with pot hoc Sidak's multiple comparisons test. Two group comparisons for the number of average cocaine infusions during the last 3 days of self-administration and beam breaks were performed using unpaired t-tests. The incubation score was calculated as WD50 - WD1 active nose pokes and assessed using Mann Whitney test. For all analyses, significance was set at $p<0.05$. The data were presented as group mean ± S.E.M.
